# Widespread employment of conserved *C. elegans* homeobox genes in neuronal identity specification

**DOI:** 10.1101/2022.04.29.490095

**Authors:** Molly B. Reilly, Tessa Tekieli, Cyril Cros, G. Robert Aguilar, James Lao, Itai Antoine Toker, Berta Vidal, Eduardo Leyva-Díaz, Abhishek Bhattacharya, Steven J. Cook, Jayson J. Smith, Ismar Kovacevic, Burcu Gulez, Robert Fernandez, Elizabeth F. Bradford, Yasmin H. Ramadan, Paschalis Kratsios, Zhirong Bao, Oliver Hobert

**Author notes:** equal contributors (Molly B. Reilly, Tessa Tekieli, Cyril Cros). Correspondence (O.H.).

## Abstract

Homeobox genes are prominent regulators of neuronal identity, but the extent to which their function has been probed in animal nervous systems remains limited. In the nematode *Caenorhabditis elegans*, each individual neuron class is defined by the expression of unique combinations of homeobox genes, prompting the question of whether each neuron class indeed requires a homeobox gene for its proper identity specification. We present here progress in addressing this question by extending previous mutant analysis of homeobox gene family members and describing multiple examples of homeobox gene function in different parts of the *C. elegans* nervous system. To probe homeobox function, we make use of a number of reporter gene tools, including a novel multicolor reporter transgene, NeuroPAL, which permits simultaneous monitoring of the execution of multiple differentiation programs throughout the entire nervous system. Using these tools, we add to the previous characterization of homeobox gene function by identifying neuronal differentiation defects for 12 homeobox genes in 20 distinct neuron classes that are mostly unrelated by location, function and lineage history. 10 of these 20 neuron classes had no homeobox gene function ascribed to them before, while in the other 10 neuron classes, we extend the combinatorial code of transcription factors required for specifying terminal differentiation programs. Furthermore, we demonstrate that in a particular lineage, homeotic identity transformations occur upon loss of a homeobox gene and we show that these transformations are the result of changes in homeobox codes. Combining the present with past analysis, 111 of the 118 neuron classes of *C. elegans* are now known to require a homeobox gene for proper execution of terminal differentiation programs. Such broad deployment indicates that homeobox function in neuronal identity specification may be an ancestral feature of animal nervous systems.

## INTRODUCTION

Nervous systems are composed of diverse sets of neuron types, each characterized by the expression of specific gene batteries that define the structural and functional features of that mature neuron type. A fundamental question in developmental neurobiology is whether there are common organizational principles for how individual neuron types acquire their unique identities. One approach to uncover such common principles is to comprehensively analyze neuronal differentiation programs throughout a given nervous system and determine whether a specific set of rules or features recur in the specification of different neuron types in different parts of the organism. We have engaged in such holistic analysis in the nematode *C. elegans*, which contains a nervous system with substantial cellular diversity but limited overall number: 302 neurons in the hermaphrodite which fall into 118 classes (White et al., 1986). In our search for common principles in *C. elegans* neuron type specification, several themes have emerged: (1) the direct, coordinated control of neuron type specific gene batteries by so-called “terminal selectors” (Glenwinkel et al., 2021; Hobert, 2008, 2016b); (2) the overrepresentation of homeodomain transcription factors as terminal selectors (Hobert, 2021; Reilly et al., 2020) and (3) the codification of individual neuron identities by distinct combinations of homeodomain proteins, unique for each individual neuron type (Hobert, 2021; Reilly et al., 2020; Vidal et al., 2022).

Since the DNA binding sites of a single transcription factor does not encode enough specificity to select downstream targets genes in vast genome sequence space, transcription factors usually operate in combination with other transcription factors (Mann et al., 2009; Reiter et al., 2017; Sousa and Flames, 2022). In the context of homeodomain proteins in the *C. elegans* nervous system, combinatorial functions of co-expressed homeodomain proteins usually been inferred genetically through removal of co-expressed homeobox genes, either in isolation or in combination, resulting in neuronal differentiation defects. In some cases, the biochemical basis for such combinatorial activity has been deduced: For example, in the case of the light touch receptor neurons, the POU and LIM homeodomain proteins UNC-86 and MEC-3 bind cooperatively to target gene promoters to determine the fully differentiated state of these neurons (Xue et al., 1993); similarly, the Prd and LIM homeodomain proteins CEH-10 and TTX-3, whose expression uniquely overlaps in the cholinergic AIY interneurons, show cooperative binding to *cis-*regulatory elements of members of the gene battery that define the terminally differentiated state of the AIY neurons (Wenick and Hobert, 2004). In other cases, homeodomain proteins appear to bind to target gene promoters independently of one another, but their joint presence is still required for target gene expression (Doitsidou et al., 2013; Zhang et al., 2014).

In this paper, we set out to further probe the extent to which homeobox genes, and combinations thereof, are involved in neuronal identity specification. First, we further refine our atlas of homeodomain protein expression throughout the nervous system and, second, we define the impact of loss-of-function alleles of 12 homeobox genes on neuronal identity specification throughout the entire nervous system. One pillar of this mutant analysis is the recently described NeuroPAL transgene, which expresses more than 40 distinct neuronal identity markers throughout the entire *C. elegans* nervous system (Yemini et al., 2021). Together with other available molecular markers we identify 12 homeobox genes involved in the specification of 20 different neuron classes (of the total of 118 *C. elegans* hermaphrodite neuron classes), 10 of which had no previously known regulator. This mutant analysis therefore expands our understanding of homeobox gene function in neuronal identity control, arguing that homeobox genes play a central and perhaps ancestral role in neuronal identity specification.

## RESULTS

### Reporter alleles refine some homeodomain protein expression profiles

Precise knowledge of the expression pattern of a gene provides a useful guide for mutant analysis. In our previous, genome-wide analysis of homeodomain protein expression, we made use of both CRISPR/Cas9-engineered reporter alleles, as well as fosmid-based reporters to assess expression patterns (Reilly et al., 2020). Fosmids are generally 30-50 kb genomic fragments, usually containing several genes up/downstream of a gene of interest and can be expected to include all *cis*-regulatory information of a tagged locus. Indeed, the expression pattern of many fosmid reporters is successfully recapitulated by CRISPR/Cas9 genome-engineered reporter alleles (Berghoff et al., 2021; Leyva-Diaz and Hobert, 2022; Vidal et al., 2022). A recently published nervous system wide scRNA transcriptome atlas, called CeNGEN (Taylor et al., 2021) is also largely congruent with an atlas of homeodomain expression profiles that was based on either fosmid-based reporters or CRISPR/Cas9-engineered reporter alleles (Reilly et al., 2020). We nevertheless set out to compare the expression of 17 newly available CRISPR/Cas9-engineered reporter alleles, generated either by us or obtained from the Du lab (Ma et al., 2021), with previously described fosmid-based reporter patterns and the scRNA CeNGEN atlas (Table 1). These reporter alleles fuse a reporter directly to the N- or C-terminus of the encoded homeodomain protein, thereby allowing the direct monitoring of protein expression. We observed largely congruent expression patterns, but also observed some differences in reporter profiles (Fig.1; Table 1). The two homeobox genes with the greatest difference in gene expression profiles between fosmid-based reporter and CRISPR/Cas9-engineered reporter allele are *vab-3,* the *C. elegans* Eyeless/PAX6 ortholog and *ceh-30,* one of the two *C. elegans* BarH1 homologs. In retrospect, the difference between the sites of expression of the *vab-3* fosmid reporter and the reporter allele is not surprising. First, the previously used *vab-3* fosmid reporter stood out for its weakness and variability in expression. Second, the fosmid-based reporter for *ceh-30* covered all intergenic, non-coding regions, but it did not over all intergenic region of the neighboring paralogue *ceh-31* with whom *ceh-30* may share *cis-*regulatory control elements (Fig.1; **Supp. Fig.S1**).

**Fig. 1:**
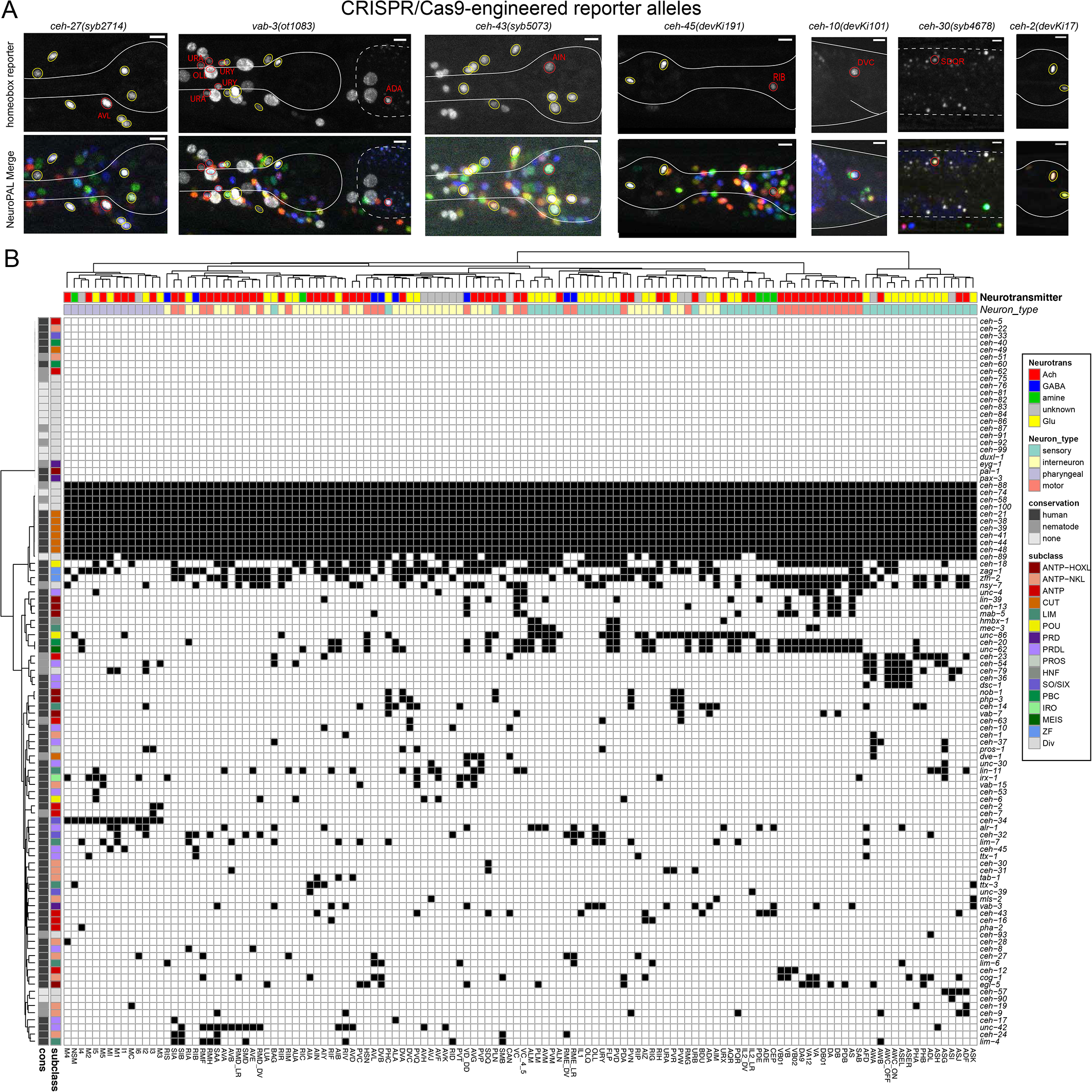
Updated expression of the homeobox gene family with reporter alleles. **A:** Representative images of homeobox reporter alleles, generated by CRISPR/Cas9 genome engineering (see strain list in **Suppl. Table S5**) with different expression than previously reported fosmid-based reporter transgenes. Neuron classes showing expression not previously noted were identified by overlap with the NeuroPAL landmark strain, and are outlined and labeled in red. Neuron types in agreement with previous reporter studies outlined in yellow. Head structures including the pharynx were outlined in white for visualization. Autofluorescence common to gut tissue is outlined with a white dashed line. An n of 10 worms were analyzed for each reporter strain. Scale in bottom or top right of the figure represents 5 um. See also **Suppl. Fig.S1** for more information on *ceh-30* and *ceh-31*. **B:** Summary of expression of all homeobox genes across the *C. elegans* nervous system, taking into account new expression patterns from panel A and all previously published data (Reilly et al., 2020). Black boxes indicate that a homeodomain transcription factor is expressed in that given neuron type and white boxes indicate that a homeodomain transcription factor is not expressed in that given neuron type. Neuron types along the x axis are clustered by transcriptomic similarity using the Jaccard index (see methods) and homeobox genes along the y axis are clustered similarly by their similar expression profiles in shared neuron types. See **Suppl. Fig.S3** for numerical representation of homeoboxes per neuron.

**Table 1:**
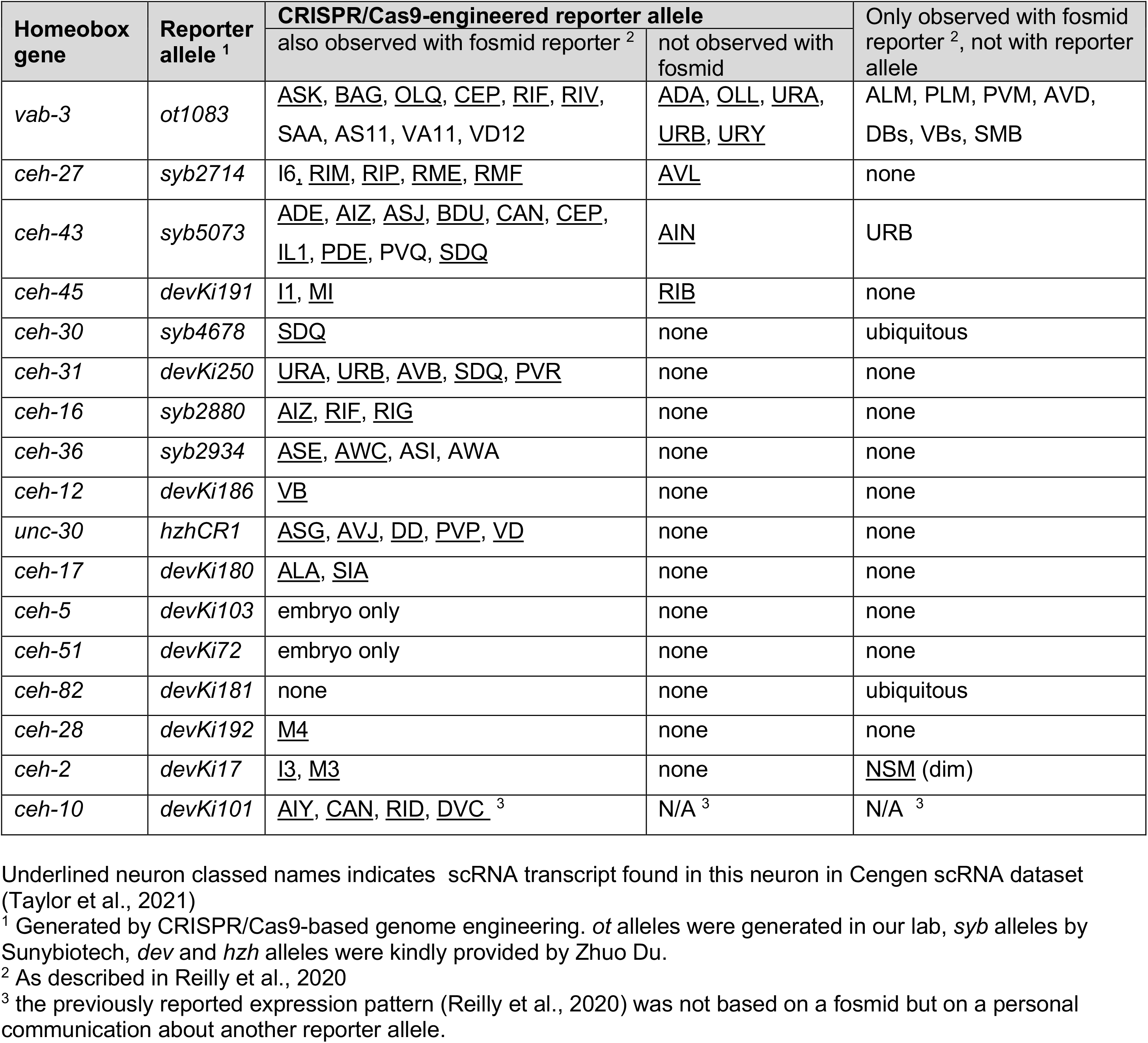
Comparing expression patterns of CRISPR/Cas9-engineered reporter alleles with fosmid-based reporter transgenes.

In two other cases, we observed that the reporter allele each added expression in one additional neuron class (Fig.1, Table 1). For example, a *ceh-45* reporter allele is expressed in I1 and MI, as previously reported with a fosmid-reporter (Reilly et al., 2020), but additional, albeit weaker expression is observed in the RIB interneurons with the reporter allele. In all these cases, the additional expression was supported by scRNA CeNGEN data (Taylor et al., 2021), but the expression level was notably lower in the respective neuron class. In these cases, the fosmid reporter may have lacked *cis*-regulatory element(s) or may have been too weakly expressed.

In one case, the EMX homolog *ceh-2,* we recapitulated previously reported I2 and M3 expression, but failed to observe CEH-2 protein expression in the NSM neuron, which had shown expression with the fosmid reporter (Reilly et al., 2020),. Since this neuron class does show scRNA transcripts of the *ceh-2* homeobox gene (Taylor et al., 2021), it is conceivable that multicopy overexpression of the fosmid reporter may have titrated out a rate-limiting posttranscriptional regulatory event.

Altogether, the revised expression patterns of homeodomain proteins further refine the nature of combinatorial homeodomain expression codes that uniquely define each neuron class (Fig.1B**; Suppl. Table S1, S2**). *In toto*, 79 of the 102 *C. elegans* homeobox genes are expressed in the nervous system, 69 of which on a neuron type-specific manner, with neuron-type specific combinatorial codes (Reilly and Hobert, 2020; Reilly et al., 2020)(this paper). We display this combinatorial coding data in a manner distinct from our previous study (Reilly et al., 2020). We first clustered neuronal cell types by similarity of scRNA-generated gene expression profiles (see Methods; **Suppl. Fig.S2**) and then mapped homeodomain expression patterns onto this matrix. This representation provides a visually tractable way to discern whether transcriptionally similar neurons tend to express the same homeodomain protein(s)(Fig.1B**; Suppl. Table S1, S2**). There are indeed several instances of such co-clustering. For example, the set of neurons that co-express the homeodomain protein CEH-34 and the set of neurons that co-express the homeodomain protein UNC-42 share more molecular similarities among themselves than with other neuron classes (Fig.1B). This is particularly notable because both the UNC-42(+) and CEH-34(+) neurons are synaptically more interconnected than expected by chance (Berghoff et al., 2021; Vidal et al., 2022).

### Mutant analysis of homeobox genes

Guided by homeodomain protein expression patterns, we sought to further implicate them in the process of terminal neuron differentiation using homeobox mutant strains. It is important to emphasize that our homeodomain protein expression profiles are entirely focused on those proteins that are continuously expressed throughout the life of a neuron and are therefore candidates to not only initiate but also maintain the differentiated state of a neuron, the original criterion for being a terminal selector of neuronal identity (Hobert, 2008, 2016b). Our mutant analysis covered neurons that fall into three categories (Table 2): (1) neurons for which no transcriptional identity regulator (terminal selector) had been described before; (2) neurons for which only a non-homeobox identity regulator had been identified; (3) neurons for which no unique functional combination of homeobox genes has been identified. The neurons that we covered in this functional analysis are functionally diverse, many have different lineage histories and are located in distinct parts of the *C. elegans* nervous system (Table 2).

**Table 2:**
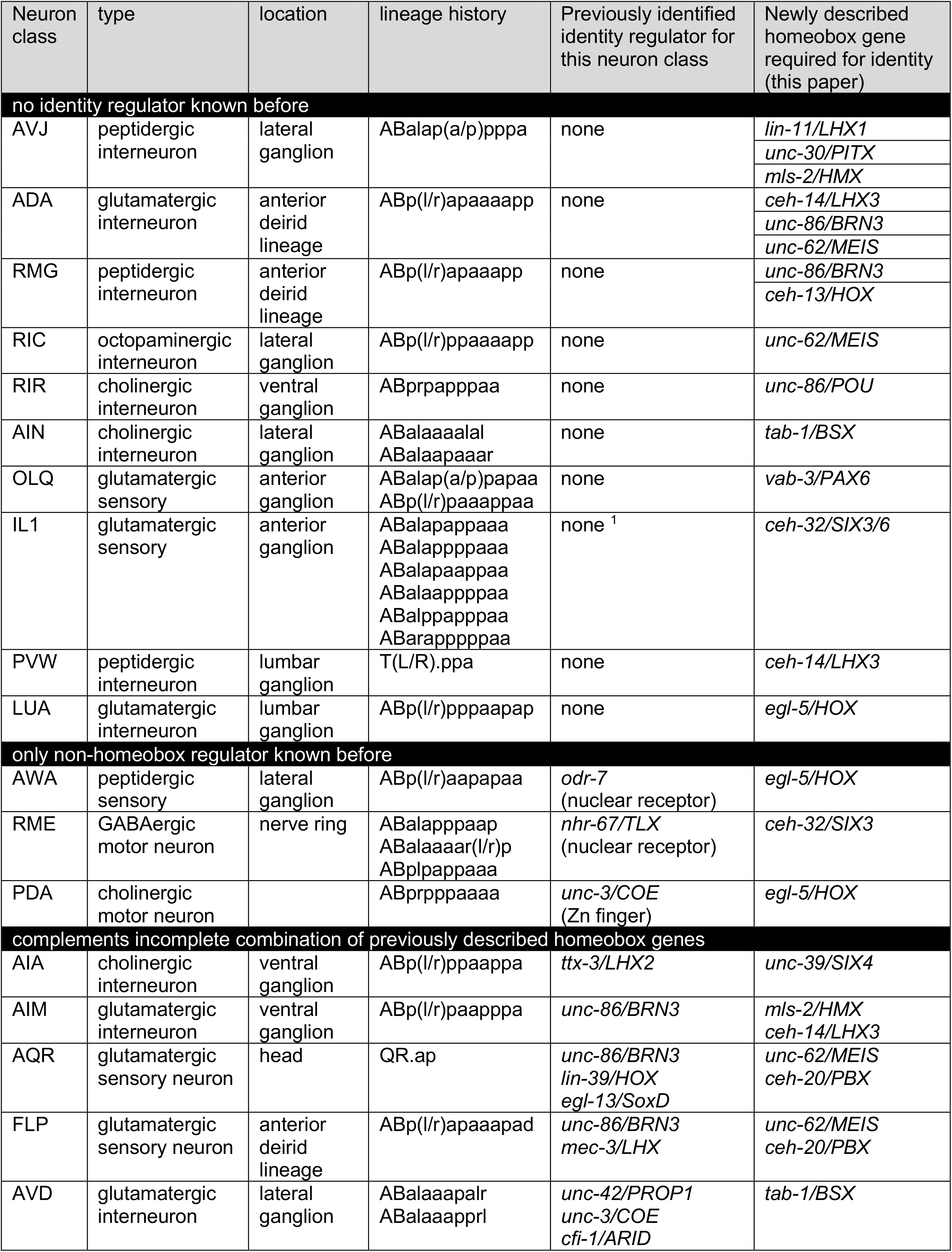
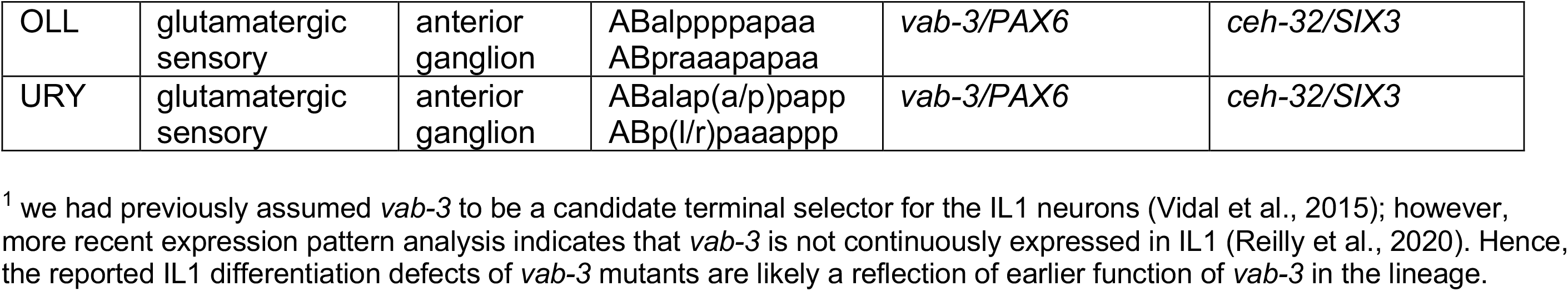
Summary of newly identified homeobox regulators of neuronal differentiation.

### The SIX-type homeobox gene *unc-39* affects differentiation of the AIA interneuron class

We have previously reported that the LIM homeobox gene *ttx-3* is required for the proper differentiation of the cholinergic AIA interneurons (Zhang et al., 2014). However, unlike in the AIY interneuron class, where TTX-3 partners with the Chx10 homolog CEH-10 (Wenick and Hobert, 2004), no homeodomain protein was shown to be a functional partner for TTX-3 in the AIA neurons. The Six4/5 homeodomain protein UNC-39 is continuously expressed in the AIA interneurons from their birth, throughout the animal’s life (Reilly et al., 2020), making it a candidate TTX-3 cofactor. The only other site of UNC-39 expression is the lateral IL2 neuron subtype (Reilly et al., 2020). We found that *unc-39(e257)* mutant animals display strong defects in the expression of six out of six tested molecular markers of AIA identity, two neuropeptides (*ins-1* and *flp-19),* the ACh vesicular transporter, *unc-17*, the choline reuptake transporter *cho-1,* a neuropeptide receptor, *dmsr-2* and a metabotropic glutamate receptor, *mgl-1* (Fig.2A), indicating that UNC-39 may indeed cooperate with TTX-3 to specify AIA identity.

**Fig. 2:**
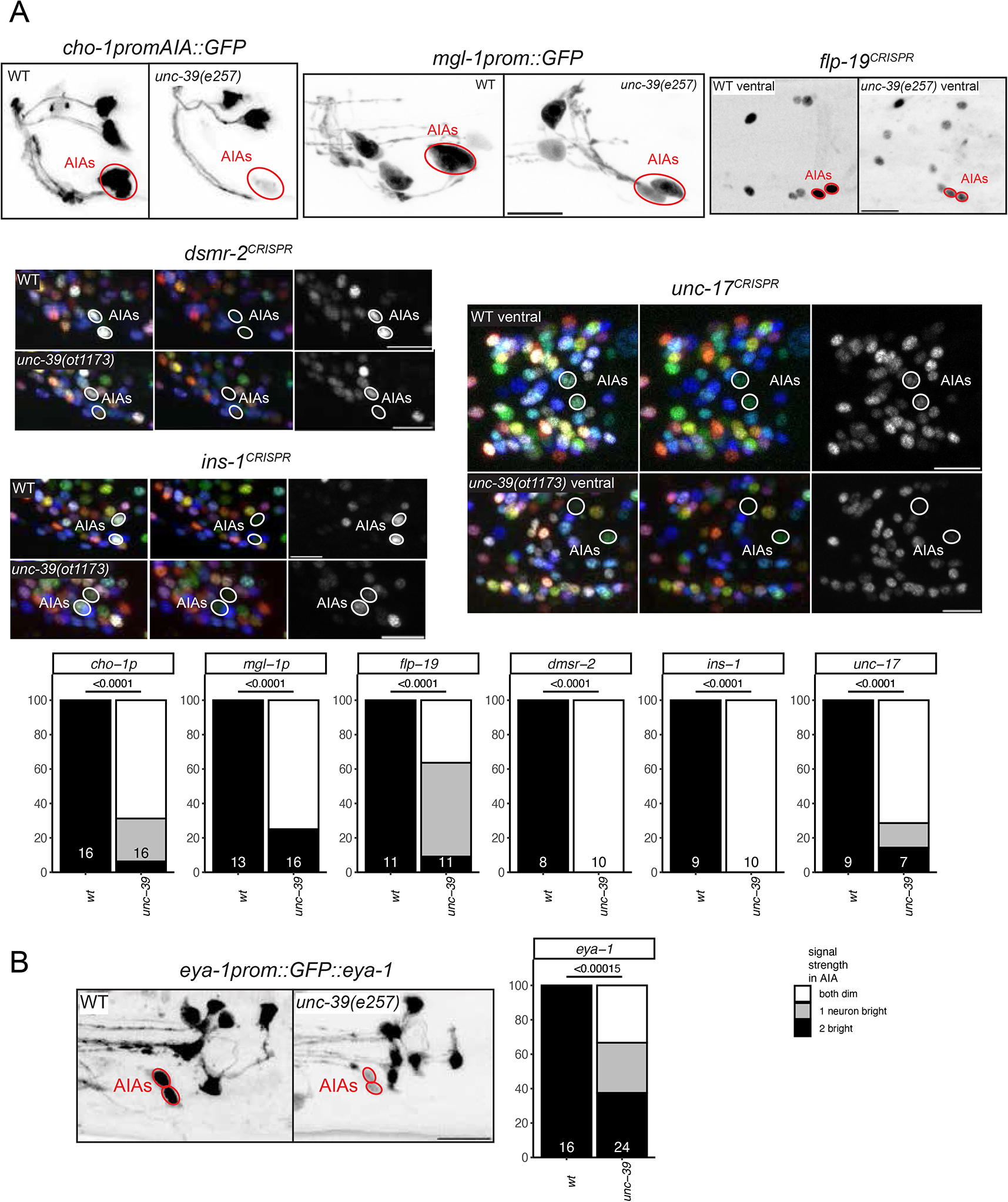
unc-39 controls differentiation of the AIA interneuron class. *unc-39*^R203Q^ mutant animals (either canonical *e257* allele or CRISPR/Cas9 genome engineered *ot1173* allele with identical nucleotide change) were analyzed. Neuron of interest is solid outlined in red or white when expressing wildtype reporter colors, and dashed red or white when one or all colors are lost. **A:** *unc-39* affects the cholinergic identity of the AIA interneuron class (*unc-17* reporter allele *syb4491* and a *cho-1* promoter fragment which is part of the *otIs653* array), and other AIA terminal identity markers: reporter alleles *dmsr-2(syb4514), ins-1(syb5452)* and *flp-19(syb3278)*, and a *mgl-1* promoter fragment *otIs327*. We did not quantify changes in AIA in the NeuroPAL color code, because it is variable in wild type. Representative images of wild type and mutant worms are shown with 10 uM scale bars. Graphs compare expression in wild type and mutant worms with the number of animals examined listed at the bottom of the bar. P-values were calculated by Fisher’s exact test. **B:** *unc-39* affects the expression of the tagged *eya-1* locus (*nIs352* transgene) in AIA.

The Eyes Absent protein is a transcriptional co-factor for many SIX domain-type homeodomain proteins across phylogeny (Amin et al., 2009; Hirose et al., 2010; Ohto et al., 1999; Patrick et al., 2013; Tadjuidje and Hegde, 2013; Vidal et al., 2022). Despite the expression of several SIX domain family members in many different parts of the *C. elegans* nervous system (Reilly et al., 2020), a genomic fragment that contains the entire, *gfp-*tagged *eya-1* locus (Furuya et al., 2005) is only expressed in pharyngeal neurons (Vidal et al., 2022) and one single extra-pharyngeal neuron class, the AIA neuron class (Fig.2B). This pattern is supported by scRNA analysis (Taylor et al., 2021)*. unc-39* is required for proper *eya-1* expression in the AIA neurons (Fig.2B). However, using two different markers (*unc-17, flp-19*), we observe no AIA differentiation defects in *eya-1* null mutants (**Suppl. Fig.S4**).

### The LIM homeobox gene *ceh-14* specifies distinct neuron classes

The PVW neuron pair is a peptidergic interneuron class located in the lumbar ganglion in the tail of the animal, with unknown function and no known identity regulator. The PVW neuron pair continuously expresses the *C. elegans* homolog of the vertebrate LHX3/4 LIM homeobox gene *ceh-14* (Cassata et al., 2000; Reilly et al., 2020). We assessed the peptidergic identity of PVW by analyzing the expression of four different neuropeptide-encoding genes that the CeNGEN scRNA atlas predicts to be expressed in PVW, *flp-21, flp-22, flp-27* and *nlp-13* (Taylor et al., 2021). To monitor *flp-22* expression, we used a previously available promoter fusion transgene (Kim and Li, 2004). For the other three neuropeptides, an SL2::gfp::H2B or T2A::3xNLS::gfp cassette was inserted at the C-terminus of the respective loci. All four reporters show expected expression in PVW (Fig.3A). In *ceh-14* null mutant animals, expression of these two of the four neuropeptide-encoding genes (*flp-22* and *flp-27*) is lost (Fig.3B).

**Fig. 3:**
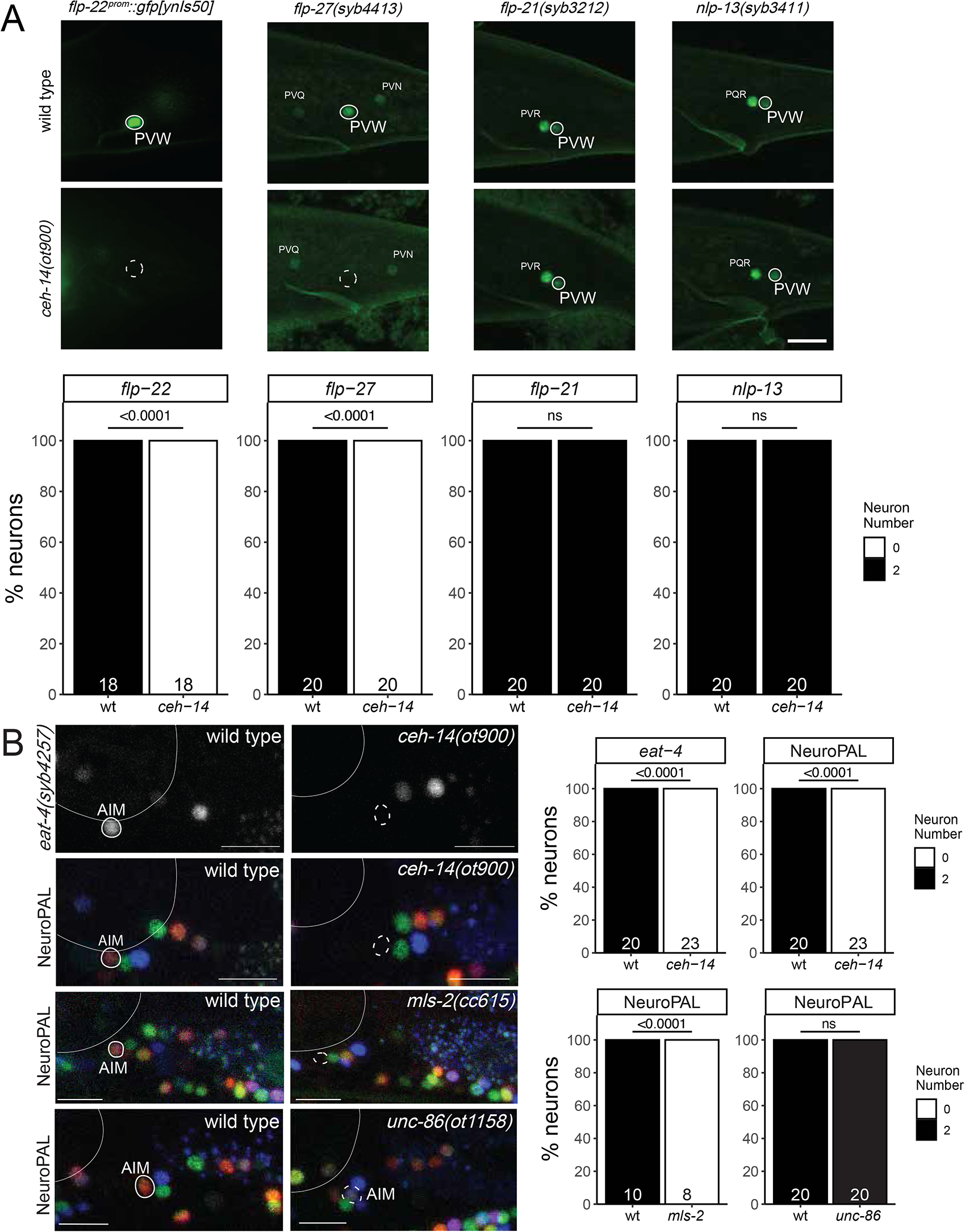
ceh-14 affect differentiation of several neuron class, in combination with different homeobox genes. **A:** *ceh-14(ot900)* mutant animals show a loss of neuropeptide-encoding genes expression in PVW, including a promoter fusion reporter transgene for *flp-22* (*ynIs50*) and a *flp-27* CRISPR reporter (*syb3213*), while expression of the neuropeptide CRISPR reporters for *flp-21* (*syb3213*) and *nlp-13* (*syb3411*) is unaffected. Neuron of interest is outlined in solid white when expressing wildtype reporter colors, and dashed white when one or all colors are lost. Representative images of wild type and mutant worms are shown with 10 μm scale bar. Graphs compare expression in wild type and mutant worms with the number of animals examined listed at the bottom of the bar. P-values were calculated by Fisher’s exact test. **B:** *ceh-14(ot900)* mutant animals show a loss of AIM marker expression, including an *eat-4* CRISPR reporter (*syb4257*) and NeuroPAL (*otIs669*) in AIM. Additionally, *unc-86(ot1158)* mutant animals show expression defects of NeuroPAL (*otIs669)* in AIM. Neuron of interest is outlined in solid white when expressing wildtype reporter colors, and dashed white when one or all colors are lost. Representative images of wild type and mutant worms are shown with 10 uM scale bars. Graphs compare expression in wild type and mutant worms with the number of animals examined listed at the bottom of the bar. P-values were calculated by Fisher’s exact test.

We also analyzed *ceh-14* function in a completely distinct set of interneurons in which CEH-14 protein is continuously expressed, the AIM neuron pair in the ventral ganglion of the head. We had previously shown that *ceh-14* affects neurotransmitter identity of AIM (Pereira et al., 2015). We confirmed these defects using an *eat-4/VGLUT* reporter allele, generated by CRISPR/Cas9 genome engineering and a previously unavailable molecular null allele of *ceh-14*, *ot900* (Fig.3B). We also analyzed expression of the NeuroPAL transgene, which expresses three markers for AIM identity, the ionotopic ACh receptor *acr-5*, the *mbr-1* transcription factor and *eat-4/VGLUT* (Yemini et al., 2021). The NeuroPAL transgene also contains a synthetic panneuronal marker, which permits assessment of the generation of a neuron. We find that in *ceh-14* null mutants, the neuron-type specific markers on the NeuroPAL transgene fail to be expressed in the AIM interneurons, while panneuronal markers are unaffected (Fig.3B). This is consistent with *ceh-14* acting as terminal selector of AIM identity.

Another homeobox gene continuously expressed in AIM is the HMX-type homeobox gene *mls-2*. *mls-2* expression overlaps with *ceh-14* exclusively in the AIM neurons. Previous work has shown that the neuropeptide-encoding gene, *flp-10,* fails to express in the AIM interneuron of *mls-2* mutants (Kim et al., 2010). To determine whether *mls-2* has similar defects as *ceh-14* mutants, we assessed expression of the NeuroPAL transgene in the AIM neurons of *mls-2* mutants. We find that *mls-2* mutants display the same color loss of the NeuroPAL transgene as *ceh-14* mutants, suggesting that *ceh-14* and *mls-2* homeobox gene collaborate to instruct AIM identity (Fig.3B).

Lastly, the BRN3 ortholog *unc-86,* a POU homeobox gene, was previously also shown to affect glutamatergic identity of the AIM neurons, as assessed by loss of *eat-4/VGLUT* fosmid reporter expression (Serrano-Saiz et al., 2013). Consistent with a role of *unc-86* in AIM differentiation, *unc-86* null mutants also display NeuroPAL color code defects (Fig.3B). In this case, however, the wild type color composition of NeuroPAL in AIM (dim *acr-5::mTagBFP*, bright *eat-4::mNeptune2.5*, dim *mbr-1::magenta*) changes from bright red to a more blue color, which corroborates loss of *eat-4/VGLUT* expression, but indicates a potential increase of expression of *acr-5* and *mbr-1* expression. *unc-86* may therefore have a more restricted impact on AIM differentiation, compared to *ceh-14* and *mls-2*.

### *mls-2/HMX, unc-30/Pitx* and *lin-11/Lhx* specify the previously uncharacterized AVJ neuron class

Apart from the AIM interneuron class, *mls-2* is also continuously expressed in the AVJ neuron pair, an interneuron class in the lateral ganglion with no presently known function or identity regulator. The AVJ interneuron class appears morphologically very similar to its neighboring AVH neuron class, and AVJ has been commonly misidentified as AVH (Cook et al., 2021). NeuroPAL provides a unique color code for AVJ, based on the expression of the neuropeptide-encoding *flp-26* gene and the AMPA Glu receptor *glr-1* (Hart et al., 1995; Yemini et al., 2021). *mls-2* affects expression of these genes, but panneuronal marker expression remains unaffected, consistent with a role of *mls-2* as a terminal selector of AVJ identity (Fig.4A). We further corroborated this notion by testing the expression of an additional neuropeptide gene, *nlp-8*, whose expression in AVJ was predicted by CeNGEN and confirmed with a reporter transgene (Fig.4B). We find proper *nlp-8* expression in AVJ to be affected by loss of *mls-2* as well (Fig.4B).

**Fig. 4:**
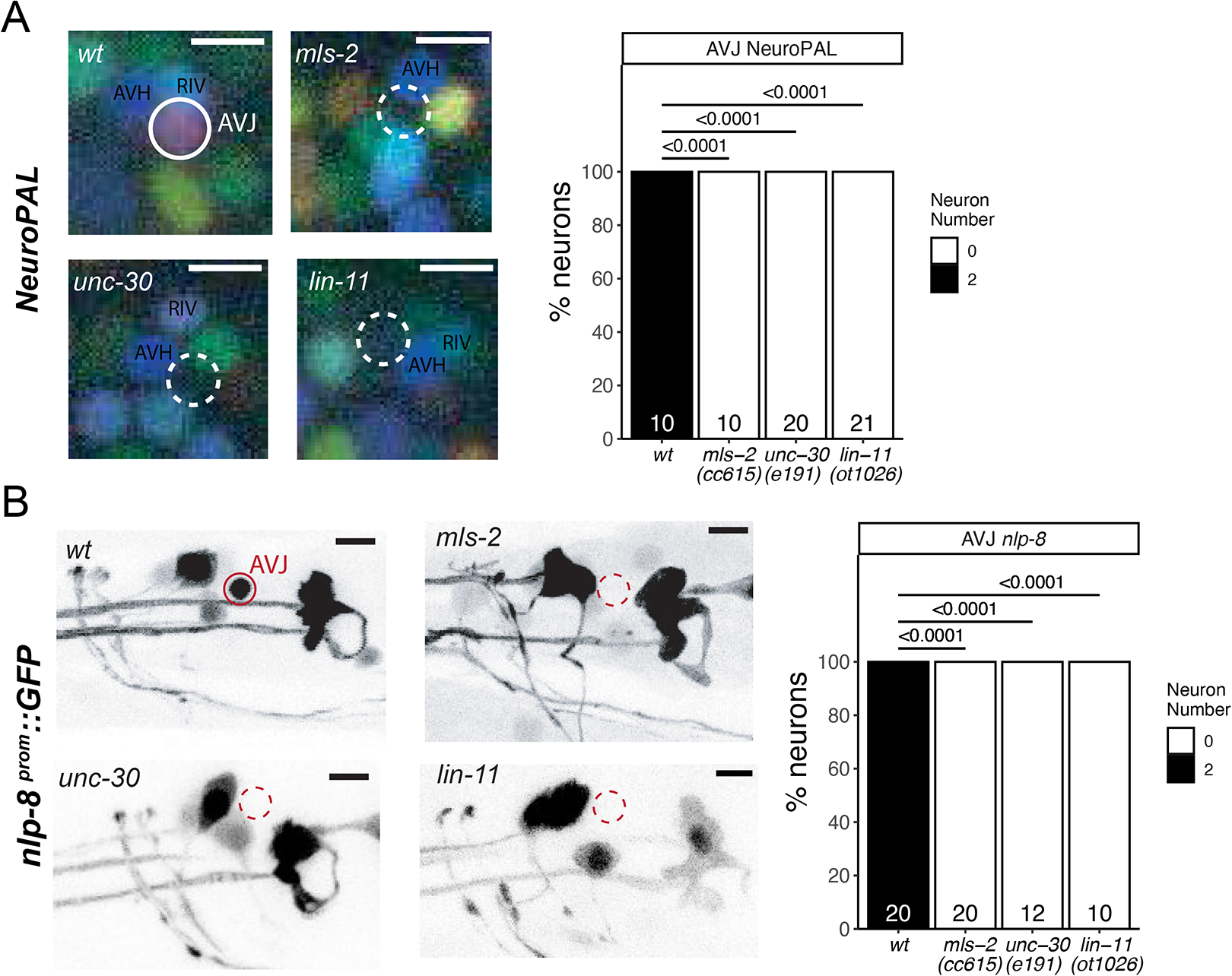
Three homeobox genes control the identity of the AVJ neuron class. **A:** *mls-2 (cc615)*, *unc-30 (e191)* and *lin-11(ot1026)* mutant animals show defects in the expression of NeuroPAL (*otIs669*) in AVJ. Neuron of interest is outlined in solid white when expressing wildtype reporter colors, and dashed white when one or all colors are lost. Representative images of wildtype and mutant worms are shown with 5 uM scale bars. Graphs compare expression in wildtype worms with the number of animals examined listed at the bottom of the bar. P-values were calculated by Fisher’s exact test. **B:** *mls-2 (cc615)*, *unc-30 (e191)* and *lin-11(ot1026)* mutant animals show defects in the expression of an nlp-8 reporter transgene (*otIs711*) in AVJ. Neuron of interest is outlined in solid red when expressing wildtype reporter colors, and dashed red when one or all colors are lost. Representative images of wildtype and mutant worms are shown with 5 uM scale bars. Graphs compare expression in wildtype worms with the number of animals examined listed at the bottom of the bar. P-values were calculated by Fisher’s exact test.

In addition to *mls-2,* the AVJ neurons co-express the LIM homeobox gene *lin-11* and the Pitx-type homeobox gene *unc-30* (Reilly et al., 2020). This homeobox code is unique for the AVJ neurons (Reilly et al., 2020). While *unc-30* and *lin-11* were previously known to control the identity of other neuron classes (Hobert, 2016a; Hutter, 2003; Jin et al., 1994), their function in AVJ has not been previously examined. Since the previously used *lin-11* mutant alleles are not unambiguous molecular nulls (Freyd et al., 1990), we generated such a null allele by deleting all coding sequences using the CRISPR/Cas9 genome engineering. Using NeuroPAL as a cell fate assessment tool, we found differentiation defects in *unc-30* and *lin-11* null mutants similar to those observed in *mls-2* mutant animals (Fig.4A). Both *unc-30* and *lin-11* also affect expression of *nlp-8* in AVJ (Fig.4B).

We also note that the AVJ gene battery shows an enrichment of phylogenetically conserved DNA binding sites of UNC-30 and MLS-2 (Glenwinkel et al., 2021), consistent with these factors acting as terminal selectors of neuron identity. LIN-11 binding sites are not specific enough to permit genome-wide analysis (Glenwinkel et al., 2021; Narasimhan et al., 2015).

### The Eyeless/Pax6 ortholog *vab-3* is required for the differentiation of several anterior ganglion neurons

As described above (Fig.1), the Pax6/Eyeless ortholog *vab-3* (Chisholm and Horvitz, 1995) is continuously expressed in a number of differentiating neurons, from their birth throughout their lifetime. *vab-3* is most prominently expressed in many neuron classes of the most anteriorly located head ganglion, called the anterior ganglion (OLQ, URY, OLL, URA, URB, CEPV and, more strongly than in all other neurons, BAG). We have previously shown that *vab-3* is required for the proper differentiation of the OLL neurons and the ventral URY neurons (URYV)(Serrano-Saiz et al., 2013), while others have shown a role in BAG neuron differentiation (Brandt et al., 2019). Seeking to extend this past analysis, we first used NeuroPAL to assess the differentiation program of anterior ganglion neurons in *vab-3* mutant mutants. We note loss of neuron-type specific NeuroPAL color codes in many anterior ganglion neurons in *vab-3* mutants, but still observe panneuronal marker expression (Fig.5A), indicating that neurons are generated, but fail to properly differentiate into specific types. Consistent with this, an *eat-4* reporter allele, expressed in all glutamatergic neurons in the anterior ganglion, shows loss of fluorescent signals in many neurons of this ganglion (Fig.5A).

**Fig. 5:**
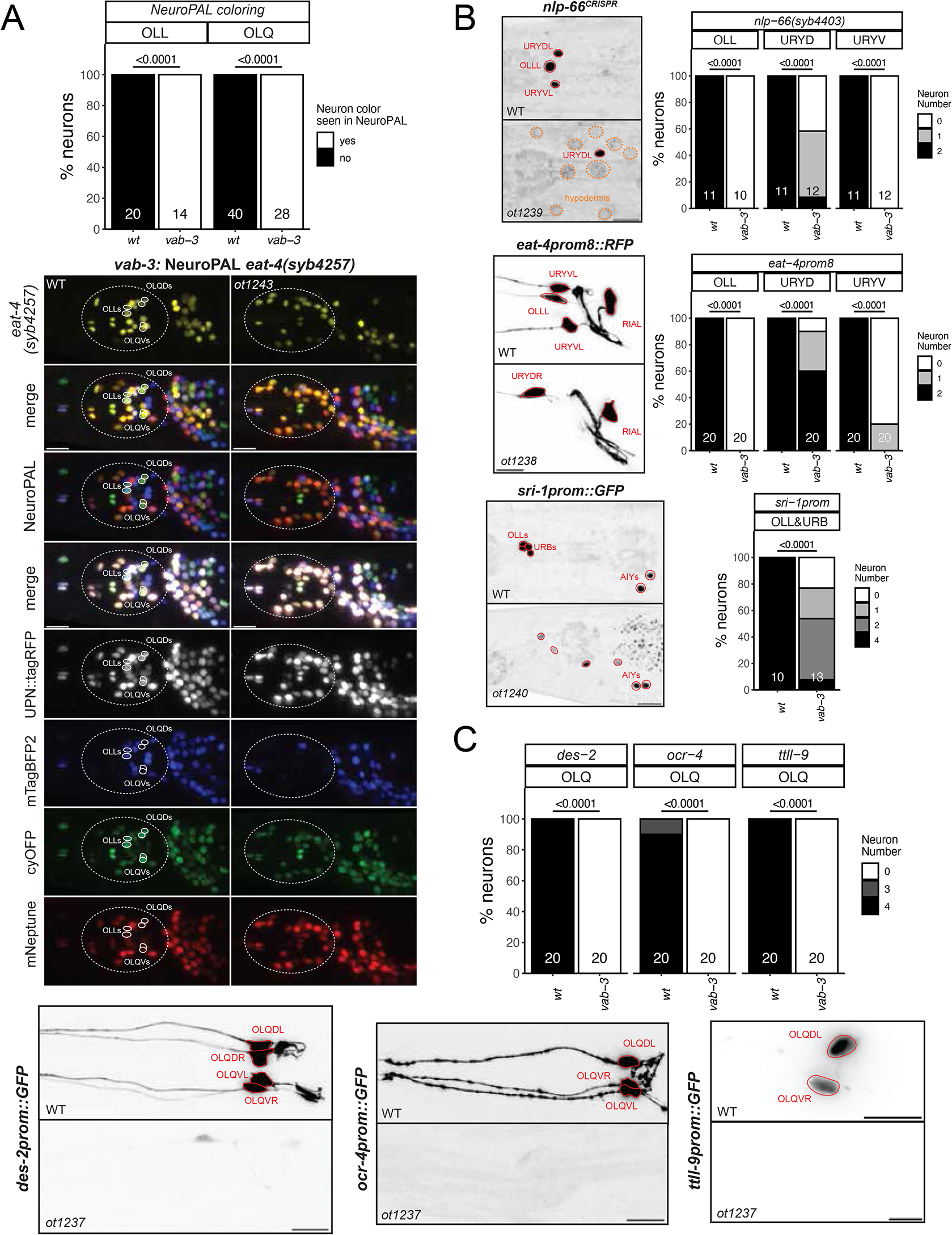
The Eyeless/Pax6 ortholog *vab-3* controls the identity of neurons in the anterior ganglion. **A:** In a *vab-3(ot12433)* mutant allele, many neurons in the anterior ganglion lose their NeuroPAL coloring (from *otIs669)* and expression of the *eat-4* reporter allele *(syb4257)*. Neurons of interest are outlined in solid white when expressing wildtype reporter colors, and dashed white when one or all colors are lost. Notably, there are much less blue neurons (URY/URA/URB but URX seem present), and the bright green OLQ and turquoise OLL are never seen. Representative images of wild type and mutant worms are shown with 10 uM scale bars. Graphs compare expression in wild type and mutant worms with the number of neurons (for n=10 WT worms / 7 *vab-3 mutant)* examined listed at the bottom of the bar. P-values were calculated by Fisher’s exact test. **B:** In *vab-3* mutant worms (*ot1239, ot1238, ot1240,* all carrying the same lesion, introduced into respective reporter background), the OLL, URY and URA markers *nlp-66(syb4403), eat-4prom8* (*otIs521*) and *sri-1 (otIs879*) are affected. Markers are often in the ventral URY than the dorsal URY. For *nlp-66*, *vab-3* mutants express *nlp-66* in hypodermis cells (circled brown). Representative images of wild type and mutant worms are shown with 10 uM scale bars. Graphs compare expression in wild type and mutant worms with the number of animals examined listed at the bottom of the bar. P-values were calculated by Fisher’s exact test. **C:** Markers of OLQ neuron identity (*ocr-4 kyEx581, ttll-9 otIs850* and *des-2 otEx7697*) are fully lost in the *vab-3(ot1237)* mutant animals. Representative images of wild type and mutant worms are shown with 10 uM scale bars. Graphs compare expression in wild type and mutant worms with the number of animals examined listed at the bottom of the bar. P-values were calculated by Fisher’s exact test.

Due to the overall disorganization of *vab-3* mutant heads, a consequence of epidermal morphogenesis defects (Cinar and Chisholm, 2004), we sought to generate and examine markers that are more specifically, if not exclusively expressed in *vab-3(+)* neurons, so that marker loss could be more unambiguously assigned to a specific neuron class. To identify such markers, we made use of the scRNA CenGEN atlas (Taylor et al., 2021). A neuropeptide encoding locus, *nlp-66*, shows highly enriched and restricted scRNA expression in URY and OLL. CRISPR/Cas9 genome engineering was used to insert an SL2::gfp::H2B reporter cassette at the 3’end of the *nlp-66* locus. As predicted, this reporter allele showed strong expression in OLL and weaker expression in both dorsal and ventral URY neuron pairs (Fig.5B). Expression of this reporter is eliminated in *vab-3* mutant animals (Fig.5B). Intriguingly, *nlp-66::gfp* reporter allele expression can be observed in what appear to be epidermal cells of *vab-3* mutant animals (Fig.5B). Brandt et al. reported a similar ectopic expression of BAG markers *ets-5* and *flp-17* in epidermal cells of *vab-3* mutants (Brandt et al., 2019).

According to the scRNA atlas (Taylor et al., 2021), as well as previous reporter gene studies (Vidal et al., 2018), the G-protein coupled receptor *sri-1* is expressed in the OLL, URB and AIY neurons. We find that a chromosomally integrated *sri-1* promoter fusion shows decreased expression in OLL and URB of *vab-3* mutant animals (Fig.5B).

The CeNGEN scRNA atlas also identified genes with very restricted expression in another *vab-3(+)* neuron class, the four radially symmetric OLQ sensory neurons. For example, transcripts for the *ttll-9* gene, encoding a tubulin tyrosine ligase-like gene, appear to be exclusive to the OLQ neurons (Taylor et al., 2021). A reporter gene fusion using 500 bp of promoter sequences confirms OLQ-exclusive expression (Fig.5C). Expression of this reporter is completely lost in *vab-3* mutant animals (Fig.5C). The *ocr-4* gene, encoding a TRP channel, was previously reported to be exclusively expressed in the OLQ neurons (Tobin et al., 2002) and we found this expression to be largely eliminated in *vab-3* mutants (Fig.5C). Lastly, in the course of dissecting the *cis*-regulatory control regions of the *des-2* locus, which encodes a degenerin-type ion channel (Yassin et al., 2001), we isolated a 1.3 kb fragment that drives exclusive reporter expression in the OLQ neurons (Fig.5C). This transgene also fails to be expressed in *vab-3* mutant animals. We conclude that *vab-3* is required for the proper execution of of most, if not all neuronal differentiation programs of the anterior ganglion in which *vab-3* is normally continuously expressed.

### The SIX3/6 ortholog *ceh-32* is also required for the differentiation of several anterior ganglion neurons

The function of PAX and SIX homeobox genes is often tightly intertwined, for example in the context of eye development (Wawersik and Maas, 2000). In *C. elegans, vab-3/PAX6* and *ceh-32/SIX3/6* function have been linked in the context of head morphogenesis (Dozier et al., 2001). The function of *ceh-32* in terminal neuron differentiation has only been reported in lateral ganglion neurons that express this gene (RIA, RMDD/V)(Cros and Hobert, 2022; Reilly et al., 2020), but not in several anterior ganglion neuron classes in which *ceh-32* and *vab-3* are co-expressed throughout the neurons’ lifetime. We find that *ceh-32* phenocopies the differentiation defects of the OLL and URY neurons observed in *vab-3* mutants. Both *nlp-66* expression, as well as expression of the glutamatergic marker *eat-4* and the tyramine receptor *ser-2* are lost in the OLL and URY neurons of *ceh-32* mutants (Fig.6A).

**Fig. 6:**
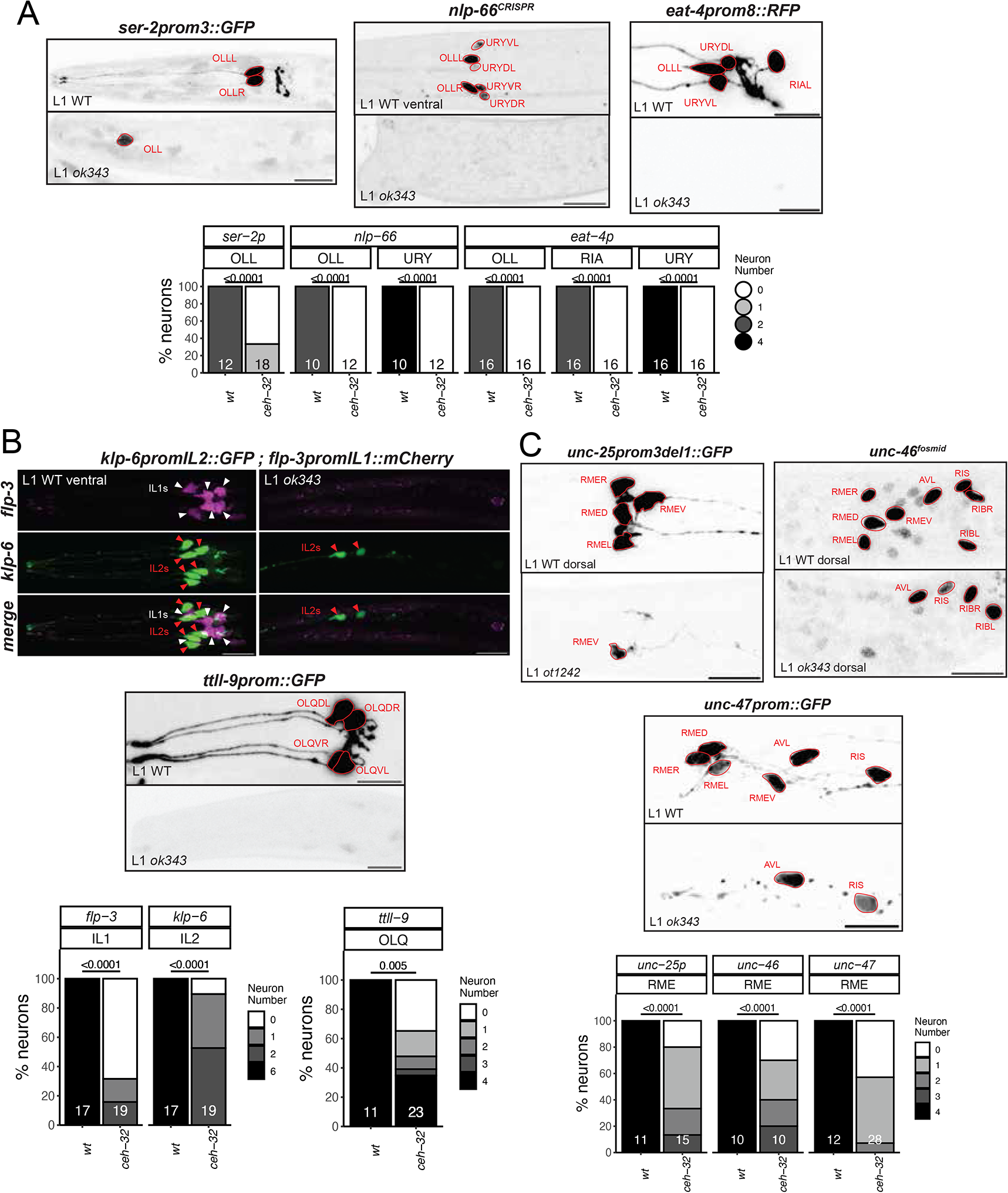
The SIX3/6 ortholog *ceh-32* controls the identity of neurons in the anterior ganglion. **A:** In *ceh-32(ok343)* null mutant animals, OLL, URY but also RIA markers are almost always lost (*ser-2prom3 otIs138, nlp-66(syb4403)*, *eat-4prom8 otIs521)*. **B:** *ceh-32* also controls IL1 identity (*flp-3* / *otIs703* marked with white triangles), and affects neurons where it is not expressed in adults (IL2: *klp-6 myIs13* marked with red triangles; OLQ: *ttll-9 otIs849*). Representative images of wild type and mutant worms are shown with 10 uM scale bars. Graphs compare expression in wild type and mutant worms with the number of animals examined listed at the bottom of the bar. P-values were calculated by Fisher’s exact test. **C:** In *ceh-32(ok343)* and *ceh-32(ot1242)* null mutant animals, the GABAergic identity of all RME classes is affected (*unc-25prom3del1 otIs837, unc-47 oxIs12*, *unc-46* fosmid *otIs568*). *ceh-32* is expressed in adults in both RME and RIB, but *ceh-32* loss did not affect RIB identity (*unc-46 otIs568* here; not shown *sto-3 otIs810*). Representative images of wild type and mutant worms are shown with 10 uM scale bars. Graphs compare expression in wild type and mutant worms with the number of animals examined listed at the bottom of the bar. P-values were calculated by Fisher’s exact test.

The glutamatergic IL1 sensory/motor neurons, another *ceh-32(+)* neuron class in the anterior ganglion, also display differentiation defects in *ceh-32* mutants. Specifically, we find that expression of the neuropeptide gene *flp-3* is lost in the IL1 neurons (Fig.6B). A similar defect has been observed in *vab-3* mutants (Vidal et al., 2015); however, since *vab-3* is not expressed throughout IL1 development (Reilly et al., 2020), its function may be restricted to earlier progenitor stages, perhaps upstream of *ceh-32.* A potential progenitor role is also evident for *ceh-32* in the OLQ and IL2 neuron classes. Both neuron classes fail to properly express differentiation markers in *ceh-32* mutants (Fig.6B). While there is no robust expression of *ceh-32* in postembryonic OLQ and IL2 neurons (Reilly et al., 2020), *ceh-32* expression is evident in the lineage that generates both OLQ and IL2 neurons (Ma et al., 2021).

Apart from the mature anterior ganglion neuron classes OLL, URY and IL1, *ceh-32* is expressed in a number of additional postmitotic neurons, including the GABAergic RME motor neurons, which are also located in the anterior ganglion and which do not express *vab-3*. A nuclear hormone receptor, *nhr-67*, but no homeobox gene was previously shown to affect the identity of all RME neurons (Gendrel et al., 2016; Sarin et al., 2009). Using three distinct markers of terminal GABAergic identity, *unc-25/GAD, unc-46/LAMP* and *unc-47/VGAT*, we find that *ceh-32* affects the identity acquisition of the RME neuron class (Fig.6C). GABAergic neuron identity was not affected in the RIB neurons, which normally also express *ceh-32* (similarly, the RIB-specific *sto-3* marker is also unaffected in the RIB neurons of *ceh-32* mutants). We conclude that *vab-3* and *ceh-32* are required for the proper terminal differentiation of a partially overlapping set of neurons in the anterior ganglion.

### *tab-1* functions in a lineage that produces the AIN and AVD neurons

The cholinergic AIN interneuron class is another neuron class for which no identity specifier has been previously identified. The AIN neuron pair is labeled with three markers by NeuroPAL (*flp-19, cho-1* and *mbr-1*) and, being cholinergic, also expresses *unc-17/VAChT* (Pereira et al., 2015). In addition, the CeNGEN scRNA atlas noted very strong and selective expression of a neuropeptide, *nlp-42*, in AIN. We tagged the *nlp-42* locus with a T2A-based reporter cassette and confirmed selective expression in AIN (Fig.7). Armed with these markers we first considered the LIM homeobox gene *ttx-3*, which is continuously expressed in AIN (Reilly et al., 2020; Zhang et al., 2014) and which is known to control the cholinergic identity of two other amphid interneuron (“AI”) classes, the AIY and AIA interneurons (Wenick and Hobert, 2004; Zhang et al., 2014). We found no obvious AIN differentiation defects in *ttx-3* mutants, as assessed with NeuroPAL, the *nlp-42* reporter allele and *unc-17* fosmid-based reporter (Fig.7A). Another homeobox gene expressed in the postmitotic AIN neurons is *tab-1,* a deeply conserved homeobox gene homologous to the *Drosophila* Brain-specific homeobox (Bsh) gene (Jones and McGinnis, 1993) and vertebrate Bsx genes (Chu and Ohtoshi, 2007). We found that NeuroPAL as well as *nlp-42* and *unc-17/VAChT* signals were absent in the AIN neurons of *tab-1* mutants (Fig.7A). The only other cholinergic neuron that expresses *tab-1* postmitotically throughout their life is the AVD command interneuron class. The NeuroPAL and *unc-17/VAChT* signals are absent in the AVD neurons of *tab-1* mutants as well (Fig.7A).

**Fig. 7:**
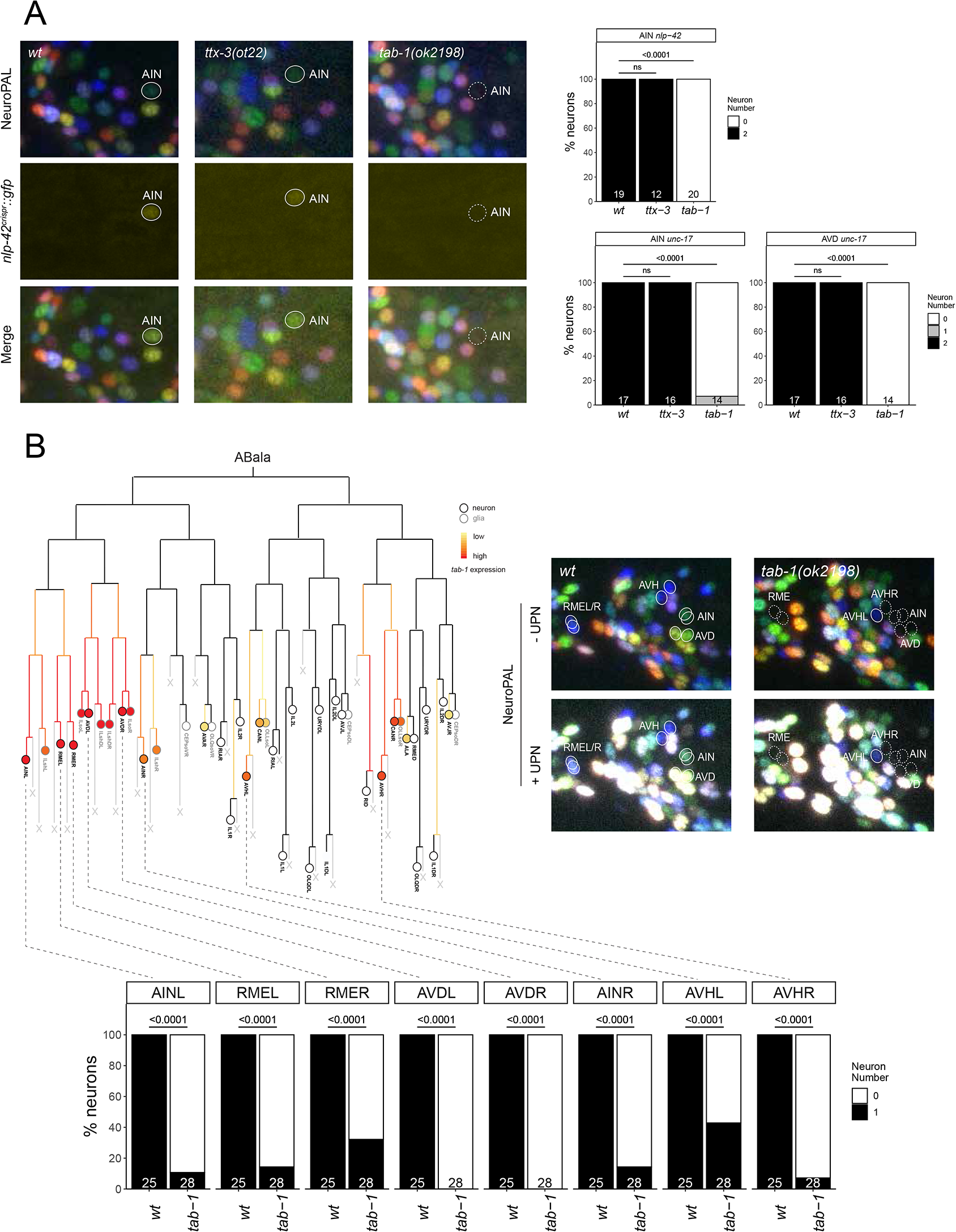
tab-1 regulates the differentiation of various neurons in the ABala lineage. **A:** In *tab-1(ok2198)* mutants, expression of both *nlp-42(syb3238)* and NeuroPAL reporters in AIN are lost. *tab-1(ok2198)* mutants also showed defects in *unc-17(otIs576)* reporter expression in AIN and AVD (image not shown). No loss of reporter expression was observed in *ttx-3(ot22)* mutants. **B:** *tab-1* is expressed in various neurons derived from the ABala lineage (adapted from Ma et al., 2021). In *tab-1(ok2198)* mutants, defects in NeuroPAL reporter expression, including ultrapanneuronal (UPN) reporter expression, are seen in neurons which express *tab-1* embryonically. In all panels, neurons of interest are outlined in solid white when expressing wildtype reporter colors, and dashed white when one or all colors are lost. P-values were calculated by Fisher’s exact test.

Notably, the panneuronal signal from the NeuroPAL transgene is also not expressed in the AIN and AVD neurons of *tab-1* mutants, suggesting a proneural role of *tab-1*. The AIN and AVD neurons are generated from the ABalaa progenitor lineage (Sulston et al., 1983)(Fig.7B). The recently reported embryonic expression of *tab-1* can be observed very early throughout the entire ABalaa progenitor lineage (Ma et al., 2021)(Fig.7B), indicating that *tab-1* may have an early role in controlling progenitor fate in this lineage. Consistent with this notion, another class of neurons generated from the ABalaa lineage, the lateral RME neuron pair also require *tab-1* for their proper differentiation (Gendrel et al., 2016)(Fig.7B). Outside the ABalaa lineage, *tab-1* is also expressed in neuroblasts that generate the AVH neuron, a neuron class that employs *unc-42* and *hlh-34* as terminal selectors (Berghoff et al., 2021). These neurons also do not acquire any NeuroPAL color code in *tab-1* mutants (Fig.7B). Since *tab-1* is not continuously expressed in AVH (Reilly et al., 2020), we again surmise an early, rather than late terminal differentiation/maintenance function of *tab-1*.

### The *egl-5* HOX gene is required for neuronal identity specification in head and tail neurons

The expression of HOX cluster genes is traditionally thought to be absent in anterior cephalic structures (Hombria et al., 2021). We were therefore intrigued to observe expression of the *egl-5* HOX cluster gene, the *C. elegans* homolog of posterior Abdominal-B-type HOX genes that normally pattern posterior structures (Chisholm, 1991; Wang et al., 1993), in the AWA olfactory sensory neuron pair in the head of the animal (Reilly et al., 2020). The identity of this neuron was previously shown to be controlled by the *odr-7* nuclear hormone receptor, which regulates the expression of the olfactory receptor ODR-10 (Sengupta and Bargmann, 1996), as well as other identity features of AWA (Glenwinkel et al., 2021). We found that in *egl-5* mutants, *odr-10* expression is also lost from AWA neurons and so is the characteristic NeuroPAL color code for AWA (Fig.8A). We instead observed ectopic expression of *odr-10* in other, positionally very distinct neuron classes in the head of *egl-5* mutants, indicating neuronal identity transformations (Fig.8A).

**Fig. 8:**
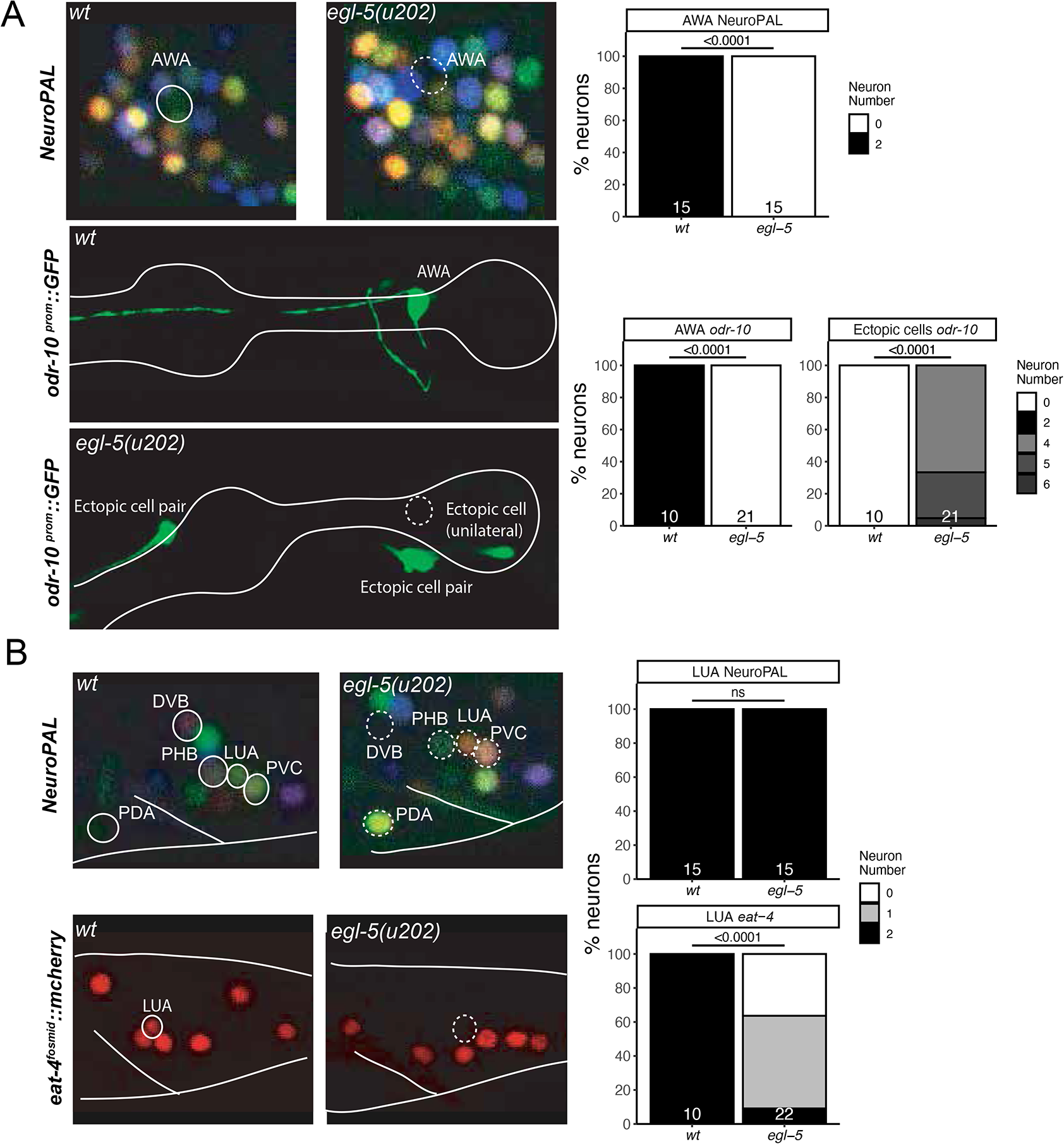
The HOX gene *egl-5* affects the differentiation of head and tail neurons. **A:** *egl-5(u202)* mutant animals show a loss of AWA marker expression, including NeuroPAL (*otIs669*) and an *odr-10* reporter transgene (*kyIs37*). Representative images of wildtype and mutant worms are shown with 5 uM scale bars. Graphs compare expression in wildtype worms with the number of animals examined listed at the bottom of the bar. P-values were calculated by Fisher’s exact test. **B:** *egl-5(u202)* mutant animals show changes in tail marker expression, including NeuroPAL (*otIs669*) in PDA, LUA, PVC and loss of eat-4 (*otIs518*) expression in LUA. Representative images of wildtype and mutant worms are shown with 5 uM scale bars. Graphs compare expression in wildtype worms with the number of animals examined listed at the bottom of the bar. P-values were calculated by Fisher’s exact test. In all panels, neurons of interest are outlined in solid white when expressing wildtype reporter colors, and dashed white when one or all colors are lost.

Since our homeodomain protein expression atlas also detected the expression of two Otx-type homeodomain proteins in the AWA neurons, *ceh-36* and *ceh-37* (Reilly et al., 2020), we tested whether those affect AWA differentiation, but found this not to be the case (**Suppl. Fig.S5**). Also, *ceh-36* and *ceh-37* displayed no apparent defects in the differentiation of the ASI neurons, another neuron class in which our homeodomain protein expression atlas found these proteins to be expressed (**Suppl. Fig.S5**).

As expected from its homology to AbdB, a posterior HOX gene, *egl-5* is also expressed in a number of neurons in the tail of the worm (Ferreira et al., 1999; Reilly et al., 2020; Wang et al., 1993) and we find that *egl-5* affects the proper specification of a number of them. Specifically, the LUA interneuron, for which no previous identity regulator was known, shows defects in NeuroPAL color coding, as well as a loss of expression of the *eat-4* reporter allele (Fig.8B). NeuroPAL color code changes were also observed in the PDA and PHB neurons, suggesting that their identity is not properly executed either (Fig.8B). PDA and PHB differentiation defects of *egl-5* mutants were also recently reported (Zheng et al., 2022), while this work was under preparation for publication. Other apparent NeuroPAL color changes in the tail of *egl-5* mutant animals are likely reflections of neuronal identity changes in the most posterior class member of ventral nerve cord motor neurons (e.g., DA9, VA12), that we previously reported with more cell-type specific markers in *egl-5* mutant (Kratsios et al., 2017) and that were corroborated by a more recent study (Zheng et al., 2022).

### The RIC neuron class requires *unc-62/MEIS* for proper differentiation

The ring interneuron (“RI”) class RIC is the only octopaminergic neuron class in *C. elegans* (Alkema et al., 2005). In addition to synthesizing octopamine (using the biosynthetic enzymes TBH-1 and TDC-1), we also found that RIC uses glutamate as a neurotransmitter. Our original mapping of the sites of *eat-4/VGLUT* expression had not identified detectable expression in RIC (Serrano-Saiz et al., 2013). However, prompted by the detection of *eat-4/VGLUT* transcripts in our scRNA brain atlas (Taylor et al., 2021), we used CRISPR/Cas9 to insert an SL2::gfp:H2B reporter cassette into the *eat-4/VGLUT* locus. This reporter allele shows the same cellular sites of expression as the previously published fosmid-based reporter, but also revealed expression in RIC (Fig.9A).

**Fig. 9:**
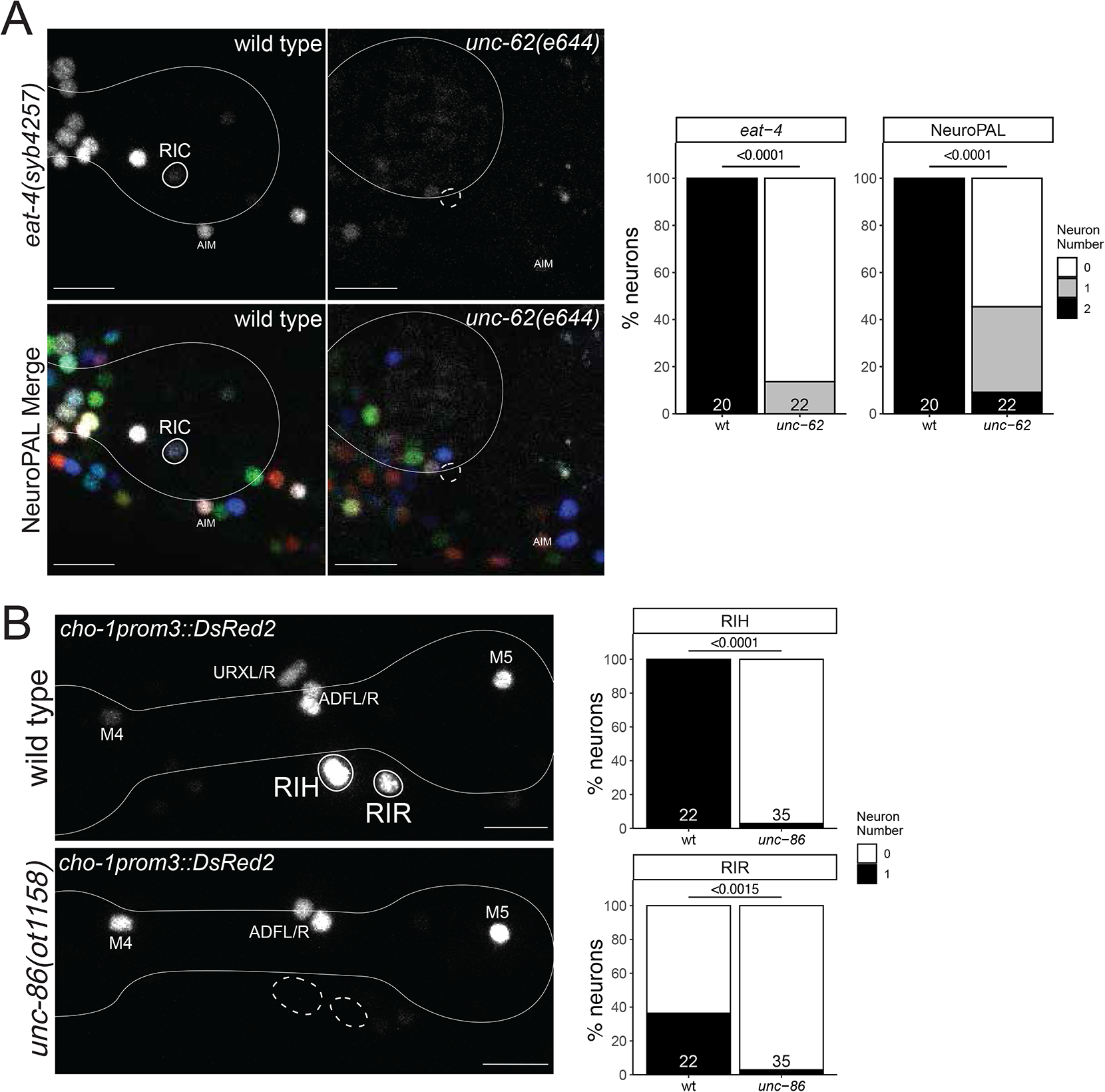
Ring interneuron (RIC, RIH, RIR) differentiation defects in homeobox gene mutants. **A:** *unc-62(e644)* mutant animals show a loss of RIC marker expression, including an *eat-4* CRISPR reporter (*syb4257)* and NeuroPAL (*otIs669*). Representative images of wild type and mutant worms are shown with 10 uM scale bars. Graphs compare expression in wild type and mutant worms with the number of animals examined listed at the bottom of the bar. P-values were calculated by Fisher’s exact test. **B:** *unc-86(ot1158)* mutant animals show loss of RIH and RIR marker expression of an extrachromosomal *cho-1prom3* reporter (*otEx4530*)(Serrano-Saiz et al., 2020). Representative images of wild type and mutant worms are shown with 10 uM scale bars. Graphs compare expression in wild type and mutant worms with the number of animals examined listed at the bottom of the bar. In all panels, neurons of interest are outlined in solid white when expressing wildtype reporter colors, and dashed white when one or all colors are lost. P-values were calculated by Fisher’s exact test.

Neuronal differentiation defects of the RIC neurons were previously reported in animals lacking two non-homeobox genes, *zip-5* and *nhr-2* (Jimeno-Martin et al., 2022), but no regulator of the homeobox family was previously known. RIC expresses the MEIS homeobox gene *unc-62* (Reilly et al., 2020). We found that in *unc-62(e257)* mutants, *eat-4::gfp* expression in RIC is lost (Fig.9A). Moreover, the fate signature of the RIC neurons in NeuroPAL (combination of *ggr-3* and *mbr-1*)(Yemini et al., 2021) is eliminated in *unc-62* mutants. Panneuronal marker expression is unaffected, indicating that *unc-62* does not control the generation, but rather the identity specification of the RIC neurons. We also note that expression of *tbh-1*, encoding a tyramine-beta hydroxylase involved in octopamine biosynthesis (Alkema et al., 2005) is not affected in *unc-62(e257)* animals, but since this allele is a non-null allele (the null allele is early embryonic lethal; (Van Auken et al., 2002)), this lack of defect is difficult to interpret.

### The cholinergic RIH and RIR interneurons require the *unc-86* homeobox gene for proper differentiation

Unlike the bilaterally symmetric RIC neuron pair, two other ring interneuron classes, the RIH and RIR neurons, are unpaired single neurons. Both neurons have a branched main process that projects into both sides of the nerve ring, but are very distinct in synaptic connectivity (White et al., 1986) and molecular profile (Taylor et al., 2021)(**Suppl. Fig.S2**). However, both neurons are cholinergic and express the POU homeobox gene *unc-86/BRN3* (Finney and Ruvkun, 1990). Previous work has shown that the serotonergic co-transmitter identity of RIH is controlled by *unc-86* (Sze et al., 2002). Cholinergic neurotransmitter identity of RIH is also affected by loss of *unc-86* (Fig.9B)(Pereira et al., 2015). Similarly, the RIR neuron, previously entirely unstudied, also fails to express a key cholinergic neurotransmitter trait (*cho-1/ChT*) in *unc-86* mutants (Fig.9B). A neuropeptide that is expressed in RIR, *nlp-52*, is, however, not affected by *unc-86* (n=12).

The RIP neurons are another ring interneuron pair that express *unc-86*. RIP is the only neuron that connects the main nervous system to the pharyngeal nervous system (Cook et al., 2020; White et al., 1986), but both its function and its developmental specification have remained unexplored. Few molecular markers existed before the advent of the scRNA CeNGEN atlas. From the RIP-specific genetic signature that CeNGEN revealed (Taylor et al., 2021), we selected two neuropeptide encoding genes that showed highly selective, strong transcript enrichment in RIP, the *nlp-51* and *nlp-73* genes. Both loci were tagged with an SL2::gfp::H2B cassette using CRISPR/Cas9 genome engineering, confirming strong and almost exclusive expression to RIP (**Suppl.Fig.S6**). However, *unc-86* null mutants displayed no defects in *nlp-51* and *nlp-73* expression in RIP (**Suppl.Fig.S6**).

We found that the Otx-type homeobox gene *ttx-1* is also expressed in RIP (Reilly et al., 2020). *ttx-1* mutant animals also displayed no *nlp-51* and *nlp-73* expression defects (**Suppl.Fig.S6**). In an *unc-86; ttx-1* double mutant strain, expression of *nlp-51* also remains unaffected (**Suppl.Fig.S6**). However, since *ttx-1* null mutants are embryonic lethal, this analysis has presently only been done with a viable *ttx-1(p767)* hypomorphic mutant and an involvement of *ttx-1* in RIP differentiation can therefore not yet be ruled out.

### Homeobox genes affect the differentiation of neurons withing the anterior deirid lineage

The glutamatergic, functionally and developmentally uncharacterized ADA neuron class is produced by the anterior deirid lineage (Fig.10A). The NeuroPAL transgene contains three molecular markers of ADA identity, *eat-4/VGLUT*, the acetylcholine receptor subunit *acr-5* and the neuropeptide *flp-26* (Yemini et al., 2021). The POU homeobox gene *unc-86/BRN3* is expressed in ADA (Finney and Ruvkun, 1990) and given the role of *unc-86/BRN3* as terminal selector in several other neurons classes (Leyva-Diaz et al., 2020)(this paper), we analyzed NeuroPAL expression in the ADA neurons of *unc-86* null mutants. We found that the NeuroPAL fate signature for ADA is lost in *unc-86* mutants (Fig.10B). Based on its continuous expression in all neurons of the anterior deirid, including ADA (Fig.10A), we considered the sole MEIS-type homeobox gene ortholog in *C. elegans*, *unc-62,* as a collaborator of UNC-86. Indeed, we observed the same NeuroPAL defects in the ADA neurons of *unc-62* mutants as we observed in *unc-86* mutants (Fig.10B). MEIS homeobox genes usually heterodimerize with PBX-type homeobox genes, of which there are three in *C. elegans* (Van Auken et al., 2002). One of the three *C. elegans* PBX genes*, ceh-20*, is co-expressed with *unc-62* in all neurons of the anterior deirid lineage, including the ADA neurons (Reilly et al., 2020)(Fig.10A). Using an *eat-4/VGLUT* fosmid reporter, we found that ADA displays differentiation defects in both *unc-62* and *ceh-20* mutant animals (Fig.10B). At least one other glutamatergic neuron class in the anterior lineage (FLP), as well as the adjacent AQR neuron also fail to acquire glutamatergic identity (i.e. *eat-4/VGLUT* expression) in *unc-62/MEIS* and *ceh-20/PBX* mutants (Fig.10B).

**Fig. 10:**
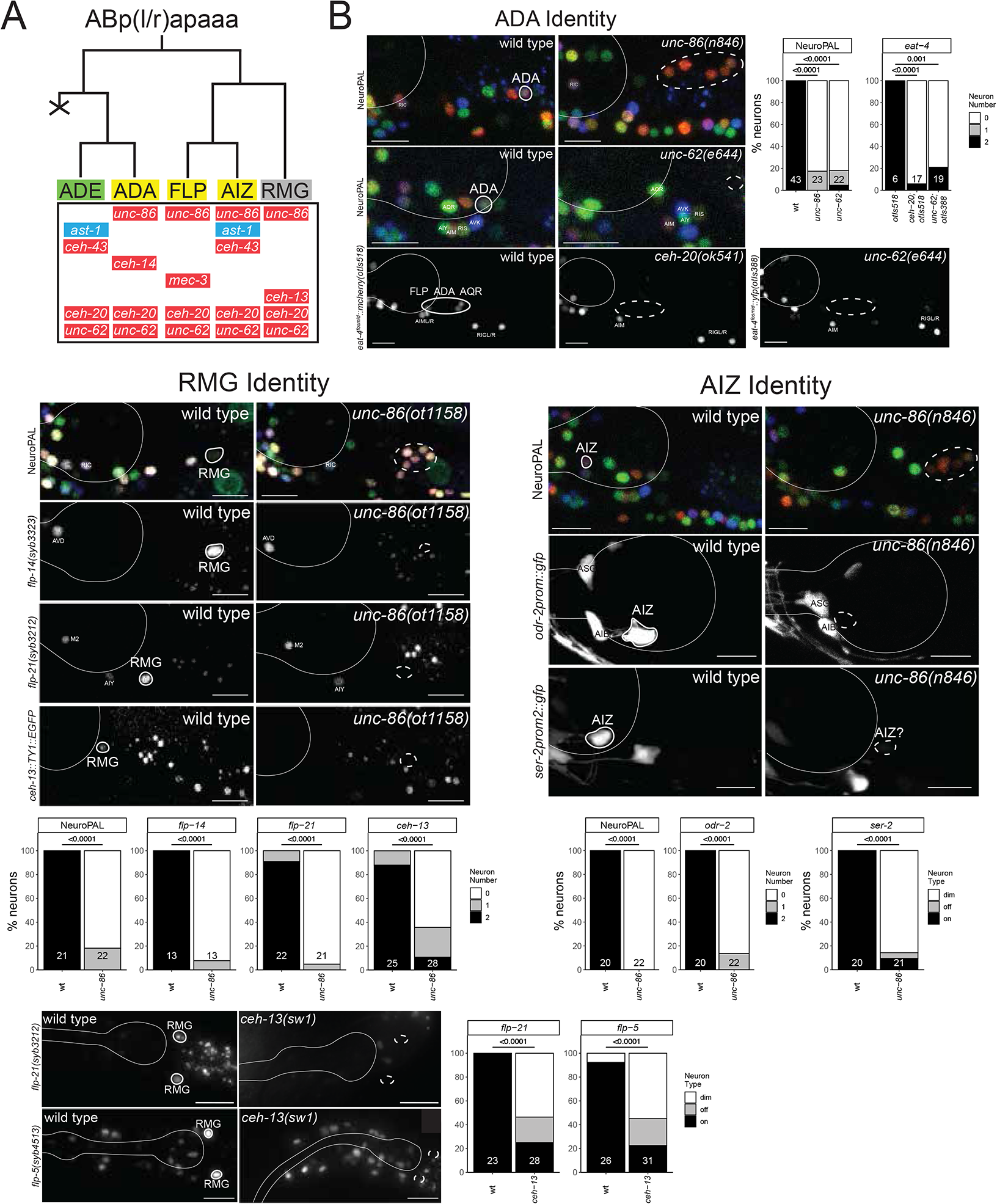
POU, MEIS and HOX genes control identity in the anterior deirid lineage. **A:** Lineage diagram depicting the generation of the anterior deirid neurons. Shown below each neuron in the lineage are the transcription factors known to be expressed in each of the respective neurons. Expression patterns for homeodomain proteins (all except AST-1) in the lineage are from (Finney et al., 1988; Reilly et al., 2020) and AST-1 is from this paper (**Suppl. Fig.S7**). Green shading: dopaminergic neuron; yellow shading: glutamatergic neuron, grey shading: peptidergic neuron, red shading: homeobox gene, blue shading: non-homeobox gene. **B:** Marker analysis in the anterior deirid lineage of wild-type and mutant animals. ADA Identity: *unc-62(e644)* and *unc-86(n846)* mutant animals show a loss of NeuroPAL (*otIs669*) in ADA. Additionally, *unc-86(n846)* mutant animals show a loss of an *eat-4* fosmid reporter (*otIs518*) in ADA. Representative images of wild type and mutant worms are shown with 10 uM scale bars. Graphs compare expression in wild type and mutant worms with the number of animals examined listed at the bottom of the bar. P-values were calculated by Fisher’s exact test. With two different *eat-4* fosmid based reporters, *unc-62* and *ceh-20* mutants also show differentiation defects in other neurons in the location of the, besides ADA (namely, FLP and AQR neurons). RMG Identity: *unc-86(ot1158)* mutant animals show a loss of RMG marker expression, including NeuroPAL (*otIs669*), a *flp-14* CRISPR reporter (*syb3323*), a *flp-21* CRISPR reporter (*syb3212*), and a *ceh-13* fosmid reporter (*wgIs756*). *ceh-13(sw1)* mutant animals show defects in the expression of two CRISPR reporters, *flp-5(syb3212)* and *flp-5(4513)*, in RMG. Representative images of wild type and mutant worms are shown with 10 uM scale bars. Graphs compare expression in wild type and mutant worms with the number of animals examined listed at the bottom of the bar. P-values were calculated by Fisher’s exact test. AIZ Identity: *unc-86(n846)* mutant animals show a loss of expression of NeuroPAL (*otIs669*) and an *odr-2* reporter transgene (*kyIs51*) and defects in the expression of a *ser-2* reporter transgene (*otIs358*) in AIZ. Representative images of wild type and mutant worms are shown with 10 uM scale bars. Graphs for NeuroPAL and *odr-2* compare expression in wild type and mutant worms with the number of animals examined listed at the bottom of the bar. Graphs for *ser-2* compare the brightness of expression (on, dim, off) in wild type and mutant worms with the number of animals examined listed at the bottom of the bar. In all panels, neurons of interest are outlined in solid white when expressing wildtype reporter colors, and dashed white when one or all colors are lost. P-values were calculated by Fisher’s exact test.

Being part of the so-called anterior deirid lineage, the ADA neuron is lineally related to another interneuron whose differentiation program was previously uncharacterized, the RMG interneuron class (Fig.10A). RMG is a so-called “hub-and-spoke” neuron that integrates signals from various sensory neurons (Macosko et al., 2009). The RMG neuron pair expresses no classic, fast acting neurotransmitter pathway machinery, i.e. is neither cholinergic, glutamatergic, GABAergic or monoaminergic, but expresses a combination of neuropeptide-encoding genes (Kim and Li, 2004; Macosko et al., 2009; Taylor et al., 2021). Like ADA, the RMG neuron class also co-expresses *unc-86* and the Meis and Pbx homologs *unc-62* and *ceh-20* (Reilly et al., 2020)(Fig.10A). Like in ADA, we find that *unc-86* is required for RMG differentiation, as assessed by its effect on neuropeptide gene expression (Fig.10B). Unlike ADA, the RMG neuron expresses the anterior most HOX cluster gene *ceh-13*, the *C. elegans* ortholog of Labial (Reilly et al., 2020)(Fig.10A), which may act to distinguish ADA from RMG. We indeed found that *ceh-13* null mutant animals fail to properly express the neuropeptides *flp-5* and *flp-21* (Fig.10B).

### *unc-86* mutants display homeotic identity transformations in the anterior deirid lineage of

Apart from ADA and RMG, other neurons within the anterior deirid lineage, specifically the AIZ and FLP neurons, also express the *unc-86/BRN3* homeobox gene (Fig.10A). Both neurons were previously shown to require *unc-86* for their proper differentiation (Serrano-Saiz et al., 2013; Topalidou and Chalfie, 2011). This notion is further corroborated with the NeuroPAL transgene; all normally UNC-86(+) neurons in this lineage fail to acquire the proper color code in *unc-86* null mutants (Fig.10B). Notably, NeuroPAL reveals that these cells instead now adopt the color that is characteristic for the non-UNC-86-expressing neuron in this lineage, the dopaminergic ADE neuron, marked with *dat-1* in the NeuroPAL transgene (Fig.10B). Upon the initial isolation of *unc-86* mutant alleles, additional cells with ectopic FIF staining, a chemical stain for dopamine, were noted in the anterior deirid lineage of *unc-86* mutant animals (Chalfie et al., 1981).

We further probed the identity of the “ectopic” NeuroPAL/FIF-positive neurons by testing five markers of the dopamine biosynthesis and vesicular packaging pathway, *bas-1/AAAD, cat-1/VMAT, cat-2/TH*, *cat-4/GTPCH*, and *dat-1/DAT.* We found all genes to be ectopically expressed throughout the anterior deirid lineage of *unc-86* mutants (Fig.11A). Another marker for ADE fate that is independent of the dopamine biosynthesis pathway is the *flp-33* neuropeptide, which we also find to be ectopically expressed throughout the anterior deirid lineage of *unc-86* mutants (Fig.11A). Similarly, the *flp-5* gene, normally expressed in ADE, FLP and RMG also becomes expressed in more cells throughout the anterior deirid lineage.

**Fig. 11:**
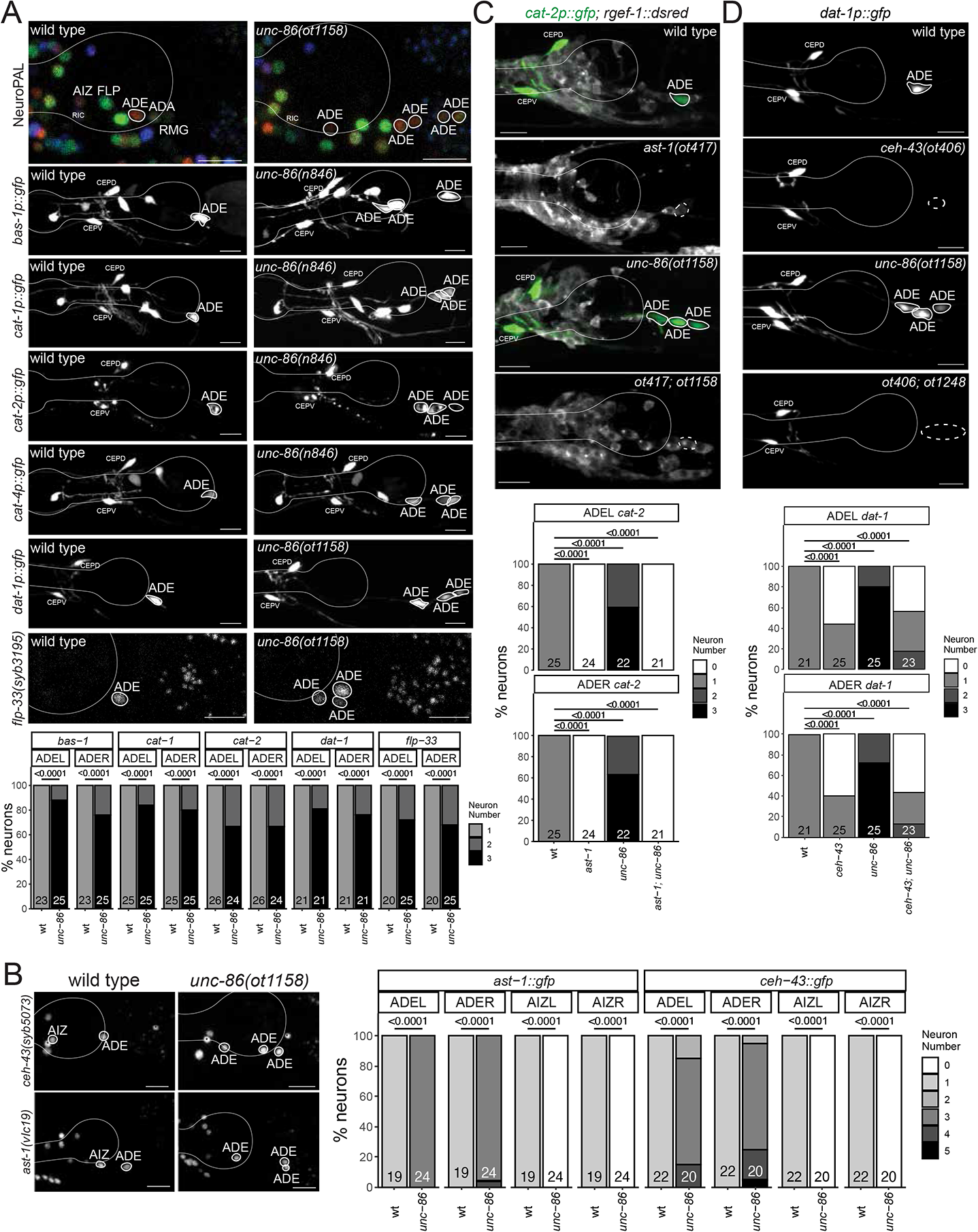
Derepression of dopaminergic terminal feature and dopaminergic regulatory signature in *unc-86* mutants. **A:** *unc-86*(*ot1158*) and *unc-86(n846)* mutant animals ectopically express markers of ADE identity, including NeuroPAL (*otIs669*), reporter transgenes for genes involved in dopamine synthesis, including *bas-1 (otIs226)*, *cat-1 (otIs224)*, *cat-2 (nIs118)*, *cat-4(otIs225)* and *dat-1(vtIs1)*, and a *flp-33* CRISPR reporter (*syb3195*). Representative images of wild type and mutant worms are shown with 10 uM scale bars. Graphs compare expression on the left side (ADEL) and the right side (ADER) in wild type and mutant worms with the number of animals examined listed at the bottom of the bar. **B:** *unc-86(ot1158)* mutant animals show a derepression of the expression in the anterior deirid of CRISPR/Cas9-engineered reporter alleles of *ceh-43(syb5073)* and *ast-1(vlc19)*. Representative images of wild type and mutant worms are shown with 10 uM scale bars (non-neuronal expression depicted with an asterisk). Graphs compare expression on the left side (ADEL and AIZL) and the right side (ADER and AIZR) in wild type and mutant worms with the number of animals examined listed at the bottom of the bar. **C,D:** *ast-1* and *ceh-43* are epistatic to *unc-86.* The derepression of expression of the *cat-2* reporter transgene (*otIs199)* in *unc-86(ot1158)* mutant is suppressed in an *ast-1(ot417)* or an *ast-1(ot406)* mutant background. Note that both alleles are hypomorphic alleles (null alleles are lethal)(Doitsidou et al., 2013; Flames and Hobert, 2009). Neurons of interest are outlined in solid white when expressing WT reporter colors, and dashed white when one or all colors are lost. Representative images of wild type and mutant worms are shown with 10 uM scale bars. Graphs compare expression on the left side (ADEL) and the right side (ADER) in wild type and mutant worms with the number of animals examined listed at the bottom of the bar. In all panels, neurons of interest are outlined in solid white when expressing wildtype reporter colors, and dashed white when one or all colors are lost and P-values were calculated by Fisher’s exact test.

We examined whether ectopic dopaminergic fate execution in the anterior deirid linage are a reflection of aberrant cell divisions leading to lineage reiterations, akin to those observed in the postdeirid lineage of *unc-86* mutants (Chalfie et al., 1981). Using 4D embryonic lineage with the StarryNite (Shah et al., 2017), we examined the cleavage patterns of six anterior deirid lineages (3 embryos) until the twofold stage in *unc-86* mutants and found no difference between these mutants and wildtype animals.

Taken together, UNC-86 not only induces specific differentiation programs in cells of the anterior deirid lineage, but also to repress an alternative neuronal identity (dopaminergic neurons) in these cells, which is normally executed by the only cell in the anterior deirid lineage that does not express *unc-86* (Fig.10A). Hence, *unc-86* appears to act as a “homeotic gene” whose loss results in neuronal identity transformations, akin to those observed in other genetic and cellular contexts in *C. elegans* (Alqadah et al., 2015; Arlotta and Hobert, 2015; Gordon and Hobert, 2015).

### Derepression of a dopaminergic regulatory code in *unc-86* mutants

What is the underlying molecular basis for homeotic identity transformation in *unc-86* mutants? The terminal differentiation of all *C. elegans* dopaminergic neurons requires a combinatorial regulatory signature composed of at least four transcription factors, the ETS domain transcription factor *ast-1*, and the three homeobox genes *ceh-43/Dlx, unc-62/Meis* and *ceh-20/Pbx* (Doitsidou et al., 2013; Flames and Hobert, 2009; Jimeno-Martin et al., 2022)(Fig.10A). *unc-62* and *ceh-20* are expressed in all neurons of the anterior deirid lineage (Fig.10A) and the data described above suggests that these factors are all required for each individual neuron fate in this lineage, apparently in combination with more restrictively expressed genes, namely *unc-86* in ADA, FLP, AIZ and RMG, and *ast-1* and *ceh-43* in ADE (Fig.10A). Consequently, in *unc-62* mutants, unlike in *unc-86* mutants, no ectopic dopaminergic fate is observed in the anterior deirid lineage (Fig.10B).

With this information in mind, an ectopic execution of dopaminergic fate throughout the anterior deirid lineage in *unc-86* mutants could mean that the complete regulatory signature for dopamine fate becomes de-repressed in *unc-86* mutants. We therefore considered the expression of both *ast-1* and *ceh-43* in the anterior lineage in wild-type and *unc-86* mutant animals. Our previous analysis of homeodomain protein expression showed that *ceh-43* is expressed in one other neuron of the anterior deirid lineage, the AIZ neuron, but it is not expressed in ADA, FLP or RMG (Reilly et al., 2020)(Fig.10A). We examined the expression of a reporter allele of *ast-1*, in which *gfp* has been inserted at the C-terminus of *ast-1* (Lloret-Fernandez et al., 2018) and found that within the anterior deirid lineage *ast-1* shows the same expression pattern as *ceh-43*. It is expressed in AIZ, but not in ADA, FLP or RMG (**Suppl. Fig.S7**; Fig.10A). The dopamine regulatory signature therefore normally exists in AIZ, but *unc-86* seemingly antagonizes the ability of *ast-1* and *ceh-43* to promote dopaminergic fate in AIZ, as inferred from AIZ normally not being dopaminergic, but turning on dopaminergic markers in an *unc-86* mutants, as described above.

Furthermore, we observed that both *ceh-43* and *ast-1* expression becomes derepressed in additional neurons in the anterior deirid lineage in *unc-86* mutants (Fig.11). To ask whether derepression of this regulatory signature can indeed be made responsible for the dopaminergic identity transformation in *unc-86* mutants, we generated *unc-86; ast-1*, as well as *unc-86; ceh-43* double mutant animals. We found that in these animals, the generation of ectopic dopaminergic neurons is suppressed (Fig.11). Hence, we conclude that in the absence of *unc-86,* a regulatory signature of dopamine fate becomes derepressed throughout the anterior deirid lineage to permit induction of dopaminergic identity. Therefore, *unc-86* normally serves to antagonize dopaminergic fate specification, either by preventing the ability of *ast-1* and *ceh-43* to promote dopaminergic fate (in AIZ) or by preventing *ast-1* and *ceh-43* expression (in other cells in that lineage).

### Several homeobox gene mutants display no apparent neuronal differentiation defects

While neuronal differentiation defects are easily apparent in many homeobox gene mutants, we also note that several neuron types display no obvious differentiation defects in single homeobox gene mutants (**Suppl. Table S3**). However, absences of defects are not straight-forward to interpret for multiple reasons: First, because our cell fate marker analysis only usually tested a small number of markers, one cannot exclude that other markers would show a defect in expression. Second, in several cases (*unc-62* and *ttx-1*) only hypomorphic alleles could be analyzed since null alleles displayed phenotypic pleiotropies that complicated the assessment of neuronal differentiation defects. Third, in several previously documented cases, even unrelated homeobox genes from different subfamilies (e.g., LIM, POU) can act redundantly in neuron identity specification (Vidal et al., 2022; Zhang et al., 2014). For example, neither *ttx-3* nor *unc-86* single mutants affect expression of several different differentiation markers of the NSM neurons, but in a *ttx-3; unc-86* double mutants these markers are completely lost (Zhang et al., 2014). A comprehensive analysis of neuronal cell fate in homeobox gene mutants may therefore require the generation of compound mutants.

## DISCUSSION

Homeobox genes have been implicated in nervous system differentiation throughout animal phylogeny (Blochlinger et al., 1988; Doe et al., 1988a; Doe et al., 1988b; Hobert, 2021; Hobert and Westphal, 2000; Holland and Takahashi, 2005; Joyner et al., 1991; Krumlauf et al., 1993; Pfaff et al., 1996; Qiu et al., 1995; Rubenstein and Puelles, 1994; Tsuchida et al., 1994). What sets apart the nematode *C. elegans* from other model systems is the extent to which homeobox genes have been implicated in neuron identity specification throughout the entire nervous system. First, expression pattern analysis of homeodomain proteins has shown that each of the 118 anatomically defined *C. elegans* neuron classes not only expresses at least one homeodomain protein, but expresses a unique combination of homeobox genes (Reilly et al., 2020). Second, homeobox genes have been shown to act as terminal selectors of neuron identity in many neuron classes of the *C. elegans* nervous system (reviewed in (Hobert, 2016a)). The first described case was the heteromeric UNC-86/MEC-3 homeodomain complex that coordinates the expression of terminal identity features of touch receptor neurons (Duggan et al., 1998; Finney et al., 1988; Xue et al., 1993). Other examples, covering many parts of the *C. elegans* nervous system, followed (reviewed in (Hobert, 2016a)). However, many neuron classes, while expressing homeobox gene combinations, had not previously been shown to require a homeobox gene for proper identity specification. Moreover, while many neuron classes were known to be affected by a specific homeobox gene, neuron-type specific homeobox co-factors had not yet been identified. For example, *unc-86* and *ceh-14* were both known to specify distinct neuron classes (Pereira et al., 2015; Serrano-Saiz et al., 2013), but it remained to be shown whether another homeobox gene dictates their ability to control distinct gene batteries in different neuron classes.

In this work, we expanded our view of homeobox gene function in the *C. elegans* nervous system. As summarized in Table 2, the present analysis has identified functions for 12 homeobox gene in 20 different neuron classes that are functionally and lineally diverse and located in different part of the nervous system. In 10 of these neuron classes no identity regulator was previously known; in 3 neuron classes only a non-homeobox gene was previously implicated in controlling terminal differentiation; and for 7 of these neuron classes homeobox genes were previously known to be involved in their differentiation, but our analysis and now added additional homeobox gene function, thereby providing unique neuron type-specific functional combinations. For example, *unc-86* and *ceh-14* function in the PHC versus the AIM neuron can now be, at least in part, explained by *unc-86* and *ceh-14* apparently cooperating with *mls-2* in the AIM neurons, but not the PHC neurons, where *mls-2* is not expressed.

Do the homeobox genes that we describe here act as master-regulatory terminal selectors? Criteria for such a role are that the protein (a) is continuously expressed during differentiation and throughout the life of the neuron to maintain the differentiated state; (b) coordinately controls the expression of many, and not just a small subset of identity features; (c) exerts this effect by direct binding to terminal effector genes. For our cell fate analysis, we often used the NeuroPAL transgene, which, depending on neuron class, contains multiple differentiation markers and/or we used a core, hardwired feature of neuronal identity, the neurotransmitter/neuropeptidergic phenotype of a neuron. Nevertheless, those markers only provide a limited view of the full differentiation program of a neuron and we have therefore not conclusively proven that the homeobox genes that we analyzed here are indeed coordinately controlling all neuronal identity aspects, to the extent that “classic” terminal selectors do.

We do note that several of the homeodomain proteins that we functionally characterized here have known and distinctive DNA binding sites (based on either ChIP and/or protein binding microarray) and those are overrepresented in the gene batteries of neurons whose identity they control (Glenwinkel et al., 2021), arguing for coordinated control of the gene battery and hence arguing for a terminal selector role of these homeodomain proteins. For example, the transcriptome of the neurons that we show in this paper to require UNC-62/MEIS for proper differentiation (ADA, FLP, RMG, AQR) show an enrichment for predicted, phylogenetically conserved binding sites of UNC-62, as determined by a phylogenetic footprinting pipeline, TargetOrtho (Glenwinkel et al., 2021). A similar enrichment of UNC-86 binding sites is observed in the *unc-86-*dependent neurons that we describe here and the same holds for MLS-2/HMX binding sites and CEH-14 binding sites in *mls-2-* or *ceh-14-*dependent neurons. Hence, we assume that in many of the cases we describe here, the respective homeodomain protein indeed serves as a terminal selector. However, we also note that in some cases, the respective homeobox gene is unlikely to act as a terminal selector. For example, the LHX3/4 ortholog affects only two out of four tested neuropeptide markers in the PVW neurons.

No matter whether each homeobox gene coordinates neuronal identity as a terminal selector or only affects specific aspects of a neuron identity, in combination with previous analysis of homeobox gene function in the *C. elegans* nervous system, we can draw a striking conclusion: most, and perhaps all, of the 118 neuron classes of the *C. elegans* hermaphrodite require at least one postmitotically expressed homeobox gene to properly execute at least some aspect of their terminal differentiation program. Specifically, according to the current tally of homeobox gene analysis (depicted in Fig.12; listed in **Suppl. Table S4**), 111 of the 118 neuron classes now have a homeobox identity regulator assigned to them. This tally takes into account not only homeobox genes acting as terminal selectors, but also homeobox genes that may “only” act as subtype selectors (e.g. the *unc-4* gene, which diversifies VA from VB motor neuron fate, in combination with the non-homeobox gene *unc-3*)(Kerk et al., 2017; Lickteig et al., 2001; Miller et al., 1992; Winnier et al., 1999).

**Fig. 12:**
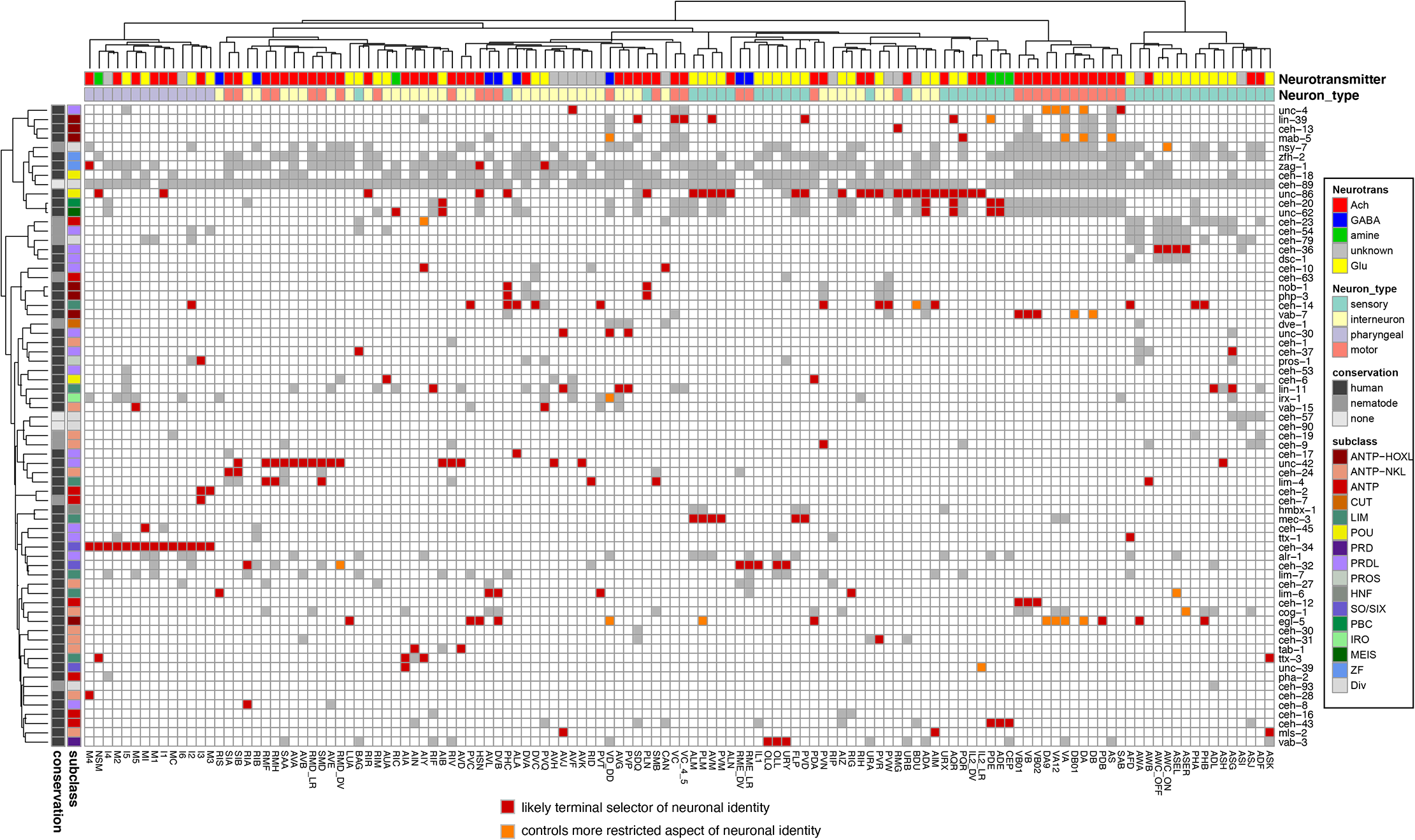
Regulation of neuron identity across the *C. elegans* nervous system by homeodomain transcription factors. Functional analysis of homeobox gene, overlayed onto the homeobox expression matrix from Fig.1B. Red boxes indicate that a homeodomain transcription factor is expressed in and likely acts as a terminal selector for a given neuron type (based on extent of functional marker analysis). Orange boxes indicate that a homeodomain transcription factor has a more restricted function in a given neuron class. Gray boxes indicate that a homeodomain transcription factor is expressed in that given neuron type, but not necessarily functionally analyzed and white boxes indicate that a homeodomain transcription factor is not expressed in that given neuron type. Panneuronal and non-neuronal homeoboxes were excluded from this representation because they do not contribute to unique neuron type codes. Neuron types along the x axis are clustered by transcriptomic similarity using the Jaccard index (see methods) and homeobox genes along the y axis are clustered similarly by their similar expression profiles in shared neuron types. See **Suppl. Table S4** for tabular list of genes and cells on which this matrix is based.

The 6 neuron classes that have not yet been found to require a homeobox gene for proper differentiation (ADF, ASI, ASJ, DVA, RIB and RIP) do express combinations of homeobox genes, but their function still requires experimental validation. An assessment of the function of several of these homeobox genes is complicated by phenotypic pleiotropies (lethality) that several of these candidate genes display.

The involvement of some homeobox genes in neuronal identity specification deserves specific note. First, HOX cluster genes can act as neuronal identity specifiers outside their more intensively studied role in ventral nerve cords, as exemplified here by *ceh-13* function in the RMG neck motor neuron or *egl-5* function in the AWA olfactory neuron. Generally, HOX gene function has been well studied in early regionalization of the nervous system, e.g. in the spinal cord (Dasen and Jessell, 2009) or the hindbrain (Parker and Krumlauf, 2020), but it is only through recent analysis in both worms (this paper)(Feng et al., 2022; Feng et al., 2019; Kratsios et al., 2017; Zheng et al., 2015a; Zheng et al., 2015b) and mice (Catela et al., 2022) that late and likely continuous roles in neuron identity specification and maintenance have become apparent. Second, the TALE-type Meis/Pbx genes, which act as common cofactors for HOX cluster genes in many different cellular contexts (Mann et al., 2009), can have HOX-independent roles. A comparison of the neuronal expression profiles of all *C. elegans* MEIS/PBX proteins with those of all HOX cluster genes has already shown that the MEIS/PBX proteins are expressed in many neuron classes that express no HOX gene (Reilly et al., 2020), but we add here functional data to this independence, showing that UNC-62 and CEH-20 acts in neurons that express no HOX cluster gene. HOX-independent roles of MEIS/PBX genes have also been observed in other organisms (Casares and Mann, 1998; Laurent et al., 2008; Remesal et al., 2020).

While in many cases, the loss of a homeobox gene results in the failure of a neuron to adopt an easily definable differentiated state, our analysis also provides a striking example of homeotic transformations in neuronal identity upon changes in homeobox codes. Specifically, we find that in the anterior deirid lineage, the POU homeobox gene *unc-86* promotes, in conjunction with neuron-type specific cofactors, the execution of several neuronal differentiation programs (FLP, ADA, AIZ, RMG), but also represses an alternative differentiation program. This alternative program – a dopaminergic neuron differentiation program – is normally promoted by two other terminal selectors, the Distalless/Dlx ortholog *ceh-43* and the ETS factor *ast-1*, but *unc-86* either represses their expression in some neurons of the anterior deirid lineage or antagonizes their function in one neuron class, the AIZ neurons, which normally express *ceh-43* and *ast-1*. One could imagine that a dopaminergic program may have been an “ancestral” state of several neurons in the anterior deirid, but the gain of *unc-86* expression in a subset of these cells diverted the differentiation programs of these cells into a different direction.

In several cases, we also infer dual roles of a homeobox gene early at progenitor stages, but also later in terminal differentiation. Such a potential dual role appears most striking for the BSH/BSX homolog *tab-1*, a homeobox gene that is already expressed early during several progenitor phases in the ABala lineage (Ma et al., 2021). Loss of *tab-1* leads to an apparent failure to generate several neurons in this lineage. However, in a subset of these neurons (AIN and AVD), *tab-1* is not only expressed during progenitor stages but is also continuously expressed during terminal differentiation throughout embryonic, larval and adult development and may be involved in initiating and maintaining the differentiated state as well. A good precedent for such a scenario is the function of the *unc-86/BRN3* POU homeobox gene in the postdeirid lineage, where *unc-86* acts at a progenitor state to prevent the progenitor from continuing to divide, but then also later in a descendant of the progenitor, the PVD harsh touch sensory neuron, to initiate and maintain its differentiated state (Chalfie et al., 1981; Finney and Ruvkun, 1990; Serrano-Saiz et al., 2018).

In conclusion, we interpret the preponderance of homeobox genes in neuronal identity control in *C. elegans* to be an indication of homeobox genes having become recruited as neuronal identity regulators early in nervous system evolution (Hobert, 2021). To further probe this issue, a systematic analysis of homeobox gene function in other organisms should be in order, but such analysis will critically depend on (a) the ability to properly assess mutant phenotypes (which, in vertebrates, may involve the experimental prevention of the execution of cell death programs often triggered after neuronal misspecification; (Serrano-Saiz et al., 2018)) and (b) the ability to conditionally target genes to avoid confounding issues such as early patterning versus late differentiation roles.

Clearly, other families of transcription factors have acquired important roles in neuronal identity specification as well, but none appear to be as broadly involved in neuronal identity specification as homeobox genes.

## ACKNOWLEDGEMENTS

We thank Chi Chen for generating transgenic animals, Zhuo Du for sending homeobox reporter alleles, Manasa Basavaraju from Marc Hammarlund’s lab for regenerating and providing an *sri-1::gfp* line, Seth Taylor in David Miller’s lab for sending and alerting us to a *vab-3* reporter allele that showed expression in tail neurons, Maryam Majeed for a transgenic reporter line, Nuria Flames for comments on the manuscript and an *ast-1* reporter allele, Ibnul Rafi for generating the *nlp-52* reporter transgene and Neda Masoudi for its initial expression analysis. Some strains were provided by the CGC, which is funded by NIH Office of Research Infrastructure Programs (P40 OD010440). This work was funded by the NIH (R01 NS039996 and R01NS110391 to O.H.; R01 NS116365-01 to P.K.; R24 OD016474 and R01 GM097576 to Z.B.; P30CA008748 to MSKCC) and the Howard Hughes Medical Institute.

## MATERIAL AND METHODS

### Strains and transgenes

A list of all strains and transgenes can be found in **Suppl. Table S5.**

### CRISPR/Cas9 genome engineering

*vab-3*, *ceh-32*, *eya-1*, *unc-86* and *lin*-*11* mutant alleles were generated using CRISPR/Cas9 and homology-directed DNA repair with a precise repair template (Dokshin et al., 2018). All these alleles are shown in **Suppl Fig.S8**. For several loci the same genomic deletion was independently generated in different reporter backgrounds, resulting in different allele names (**Suppl. Fig.S8**).

Reporter alleles were generated by Sunybiotech, inserting at the 3’end of the respective loci either a *gfp* tag (*ceh-43),* an T2A::gfp::H2B cassette (*eat-4*), or, for neuropeptide-encoding genes, an SL2::gfp::H2B cassette or an T2A::3xNLS::gfp cassette. See **Suppl. Table S5** for each individual case. We generally found T2A::3xNLS::gfp to not give as bright a signal as other ensuing SL2::gfp::H2B cassette. We will report on systematic comparison between these cassettes elsewhere.

### Clustering of neuron types by transcriptomes

To cluster neurons by transcriptomic similarities (**Fig.S2,** Fig.1, Fig.12), we used the Jaccard index distance metric and clustered those distances using the Ward.D2 clustering method in R. The Jaccard distance metric is a binary determination of similarity between two groups. In this case, two neuron groups were assessed by the number of shared genes they express divided by the shared and unshared genes in those neuron groups. Thus, this analysis does not correlate levels of gene expression, but rather treats neuron types that express the same gene as similar. We then clustered those neurons based on their Jaccard distance using the Ward.D2 clustering method, which is agglomerative clustering mechanism where every neuron type starts in a separate cluster and is combined with similar neurons such that the internal cluster variance is minimized, until all neurons are part of a single cluster. We used the hclust() function in R to perform this analysis and the relationship of all neuron types is shown as a dendrogram.

During this analysis we noted that because each neuron type was sequenced at different depths there was an inconsistent number of transcripts picked up in each neuron type. Our initial clusterings using the complete transcriptome only showed relationships that reflected the depth of sequencing of each neuron type. To rectify this, we limited our transcriptomic analysis to only the top 500 expressed genes in each neuron type, assuming that those top genes were the important transcriptomic signature of that given neuron type. Therefore, our final clustering of the neuron types is based only on the 500 most highly expressed genes in each neuron type and whether or not those genes are similar to another neuron’s top 500 genes (**Suppl. Fig.S2)**.

### Microscopy

Worms were anesthetized using 100mM of sodium azide (NaN_3_) and mounted on 5% agarose pad on glass slides. Images were acquired using confocal laser scanning microscopes (Zeiss LSM800 and LSM880) and processed using the ImageJ software (Schindelin et al., 2012). For expression of reporters, representative maximum intensity projections are shown for GFP channel as gray scale and gamma and histogram were adjusted for visibility. For mutant functional analysis, representative maximum intensity projections are shown as inverted gray scale. NeuroPAL images provided in supplement are pseudocolored in accord with ^28^. All reporter reagents and mutants were imaged at 40x using fosmid or CRISPR reagents unless otherwise noted.

### Mutant analysis scoring and statistics

Reporter expression was scored as an all-or-nothing phenotype per neuron, with expression in 0,1 or 2 neurons. Scoring data was processed in R and converted as number of expressing neurons by genotype contingency tables. Statistical analysis was then done using Fisher’s exact test under the two-sided null hypothesis. The resulting adjusted p-values are shown as less than 0.0001 when appropriate. No statistical methods were used to determine sample size prior to experiment. Based on the common standard in the field, we aimed for n greater than or equal to 10 per genotype.

### Examination of expression reagents and neuron identification

For all genes where difference of expression was noted from previously reported, we crossed the CRISPR/Cas9 generated reporter strain with the NeuroPAL landmark strain (*otIs669* or *otIs696*) and determined neuronal expression by colocalization of landmark colors and position of the neurons.

### DNA constructs

Promoter fusions for *des-2* and *ttll-9* were Gibson-cloned into the pPD95.75 backbone, using either the SphI/XmaI multiple cloning sites. A PCR fusion reporter was generated for *sri-1* following (Hobert, 2002) by Manasa Basavaraju. Primer sequences are shown below:

*des-2:* 1.3kb promoter: **fwd** gttggctgacagatcaggtg **rev** cctgtagtaaaagtaaatgtgtgttgtgtg

*ttll-9:* 500bp promoter: **fwd** ACCAAGTTCGCTTATCAGTTG **rev** cacaacaaaaaaaatccaaaaactagtcg

*sri-1:* 800bp promoter: **fwd** Gaaaattgcattatattaatgttgttcaag **rev** gagtttagcatactaaaaaATG

## SUPPLEMENTARY FIGURE LEGENDS

**Suppl. Fig.S1: Chromosomal location, reporters and summary of expression patterns of the two *C. elegans* BarH1 homologs, *ceh-30* and *ceh-31*.**

**A:** Comparison of expression of fosmid-based reporters (from Reilly et al., 2020) to CRISPR/Cas9-engineered reporter alleles (see Fig.1 for expression pattern). The ubiquitous expression of the original *ceh-30* fosmid reporter may have been an artefact. SDQ expression of the *ceh-30* reporter allele *syb4678* may be a result of cis-regulatory elements upstream of *ceh-31*, which were not included in the *ceh-30* fosmid reporter, but were included in the *ceh-31* fosmid reporter (which produces an expression pattern identical to that of the reporter allele).

**B:** Transcript abundance of *ceh-30* and *ceh-31* from the CeNGEN project, at 4 different threshold values (Taylor et al., 2021). *ceh-31* transcripts completely match the sites of fosmid reporter (*wg370)* and reporter allele expression (*devKi250)*. *ceh-30* transcripts are indeed most strongly enriched in SDQ (the only neuron class where the reporter allele *syb4678* shows expression), but is observed in other neurons as well. While those do not reporter allele expression, it is notable that they partially overlap with the sites of *ceh-31* expression. Perhaps these transcripts are produced from similar regulatory elements as the *ceh-31* transcripts, but are somehow not translated into protein.

**Suppl. Fig.S2: Clustering of neuronal cell types based on molecular similarities.**

**Suppl. Fig.S3: Numerical representation of homeobox expression data.** This data uses the expression data from **Suppl. Table S1 and S2.**

**Suppl. Fig.S4: *eya-1* null mutant does not affect expression of AIA differentiation markers.** We introduced the same CRISPR null deletion as in (Vidal 2022) in the *unc-17* and *flp-19* reporter alleles *syb4491* and *syb3278,* yielding alleles *ot1208* and *ot1209*. A small Z-stack (6-7 images at .8 micron) around the middle of the worm are shown with 10 uM scale bars. Graphs compare expression in wild type and mutant worms with the number of animals examined listed at the bottom of the bar. P-values were calculated by Fisher’s exact test.

**Suppl. Fig.S5: Effect of loss of *ceh-36* and *ceh-37* on ASI, AWA, ASE markers.**

**A:** ASI and AWA analysis in *ceh-36(gj2127)* and *ceh-37(ok642)* mutant animals. Markers used are CRISPR/Cas9-engineered reporter alleles for *ins-3(syb5421)*, *ins-6(syb5463)*, *ins-24(syb5447)*, *ins-30(syb5526)*, *nlp-2(syb5697)* and the *odr-10(kyIs37)* reporter transgene. Representative images of wild type and mutant worms are shown. Graphs compare expression in wild type and mutant worms with the number of animals examined listed at the bottom of the bar. P-values were calculated by Fisher’s exact test.

**B:** ASE analysis in *ceh-36(gj2127)* mutant animals. Markers used are CRISPR/Cas9-engineered reporter alleles for *ins-3(syb5421)*, *ins-30(syb5526)* and the *flp-6(ynIs67)* and *flp-13(ynIs37)* reporter transgenes. Representative images of wild type and mutant worms are shown. Graphs compare expression in wild type and mutant worms with the number of animals examined listed at the bottom of the bar. P-values were calculated by Fisher’s exact test.

**Suppl. Fig.S6: RIP differentiation in *unc-86* and *ttx-1* mutants.**

*unc-86(ot1158)* and *ttx-1(p767)* mutant animals don’t show loss of neuropeptide gene reporters in the RIP neuron. Representative images of *nlp-51(syb2805))* and *nlp-73(syb4406)* reporter alleles. The *nlp-51* reporter is brightly expressed in RIP and dimly expressed in AIM. The *nlp-73* reporter is expressed in the identical head neurons, but both at a moderate brightness (both reporters confirmed by NeuroPAL analysis as predicted by CENGEN scRNA data). Outside the head, *nlp-51* expresses dimly only in the tail neuron PVN beginning at the L4 stage (NeuroPAL and CENGEN scRNA data validation). The *nlp-73* reporter also expresses at a similar moderate brightness in the uv1, VC4/5, and LUA cells in addition to RIP and AIM. Mutant analysis revealed that AIM expression, but not RIP expression, is lost in *unc-86(ot1158)* animals. Expression is not impacted in any other cell types. Graphs compare RIP expression in wild type and mutant worms with the number of animals examined listed at the bottom of the bar. P-values were calculated by Fisher’s exact test.

**Suppl. Fig.S7: Expression pattern of *ast-1(vlc19)* reporter allele.** ast-1 CRISPR reporter (vlc19) is expressed in the following head neuron classes: ADE, AIN, AIZ, ASG, AVG, CEPD, CEPV, I4, I5, M3, M5, RIV, RMD, RMDD, RMDV, SMBD, SMBV, SMDD, and SMDV. Expression in the midbody, ventral nerve cord, and tail was not examined.

**Suppl. Fig.S8: Graphical representation of the homeobox mutant alleles that we generated by CRISPR/Cas9 genome engineering.** Deletions were generated by CRISPR/Cas9 genome engineering using an oligo-repair template. Identical deletions introduced into different reporter strain backgrounds get separate allele names. For *lin-11*, the deletions *ot1025* and *ot1026* are the same, but *ot1241* is different by 4 nucleotides, even though the same oligo-mediated repair template was used.

## SUPPLEMENTARY TABLE LEGENDS

**Suppl. Table S1: Homeodomain regulatory map, organized by homeodomain protein.**

This is an updated version of a table from (Reilly et al., 2020).

**Suppl. Table S2: Homeodomain regulatory map, organized by expressing neuron**. This is an updated version of a table from (Reilly et al., 2020).

**Suppl. Table S3: Homeobox mutants showing no differentiation defects in specific neuron classes.**

**Suppl. Table S4: A regulatory map of transcription factors with a role in terminal neuron differentiation**. The basis for this data is taken mainly from (Hobert, 2016a) and supplemented with data from this paper, as well as others, including (Berghoff et al., 2021; Gendrel et al., 2016; Jimeno-Martin et al., 2022; Lloret-Fernandez et al., 2018; Maicas et al., 2021; Reilly et al., 2020; Vidal et al., 2022; Zheng et al., 2022). Criteria to be included in this list is that the transcription factor is expressed throughout embryonic and postembryonic development of the respective neuron type, i.e. is likely involved not only in initiation, but also maintenance of terminal differentiation programs and the existence of mutant data that support a role in controlling marker genes. The homeobox part of this table is graphically presented in Fig.12.

**Suppl. Table S5: List of strains used in this paper.**

