## Supplementary Figures for "Widespread employment of conserved *C. elegans* homeobox genes in neuronal identity specification"

A

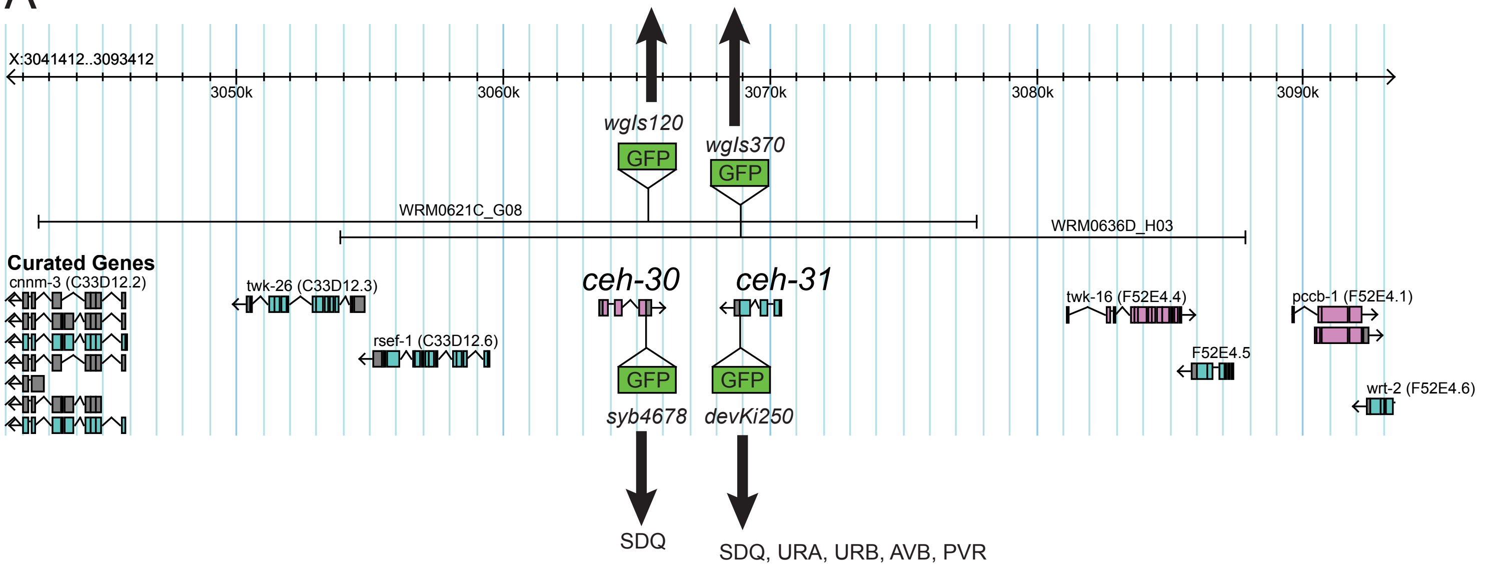


B

*ceh-30*

**CeNGEN scRNA threshold**

| 1 | | 2 | | 3 | | 4 | |
|---|---|---|---|---|---|---|---|
| SDQ | 565.303 | SDQ | 565.303 | SDQ | 565.303 | SDQ | 565.303 |
| BDU | 477.161 | BDU | 477.161 | BDU | 477.161 | BDU | 477.161 |
| PVR | 476.42 | PVR | 476.42 | PVR | 476.42 | PVR | 476.42 |
| CAN | 210.146 | CAN | 210.146 | CAN | 210.146 | CAN | 210.146 |
| RMG | 67.19 | RMG | 67.19 | RMG | 67.19 | RMG | 67.19 |
| URB | 62.236 | URB | 62.236 | | | | |
| AIN | 32.094 | AIN | 32.094 | | | | |
| AVH | 6.473 | | | | | | |
| RID | 2.954 | | | | | | |

*ceh-31*

**CeNGEN scRNA threshold**

| 1 | | 2 | | 3 | | 4 | |
|---|---|---|---|---|---|---|---|
| PVR | 6078.299 | PVR | 6078.299 | PVR | 6078.299 | PVR | 6078.299 |
| URA | 4169.736 | URA | 4169.736 | URA | 4169.736 | URA | 4169.736 |
| AVB | 3008.104 | AVB | 3008.104 | AVB | 3008.104 | AVB | 3008.104 |
| SDQ | 2426.649 | SDQ | 2426.649 | SDQ | 2426.649 | SDQ | 2426.649 |
| URB | 2036.383 | URB | 2036.383 | URB | 2036.383 | URB | 2036.383 |

Supp. Fig.S1

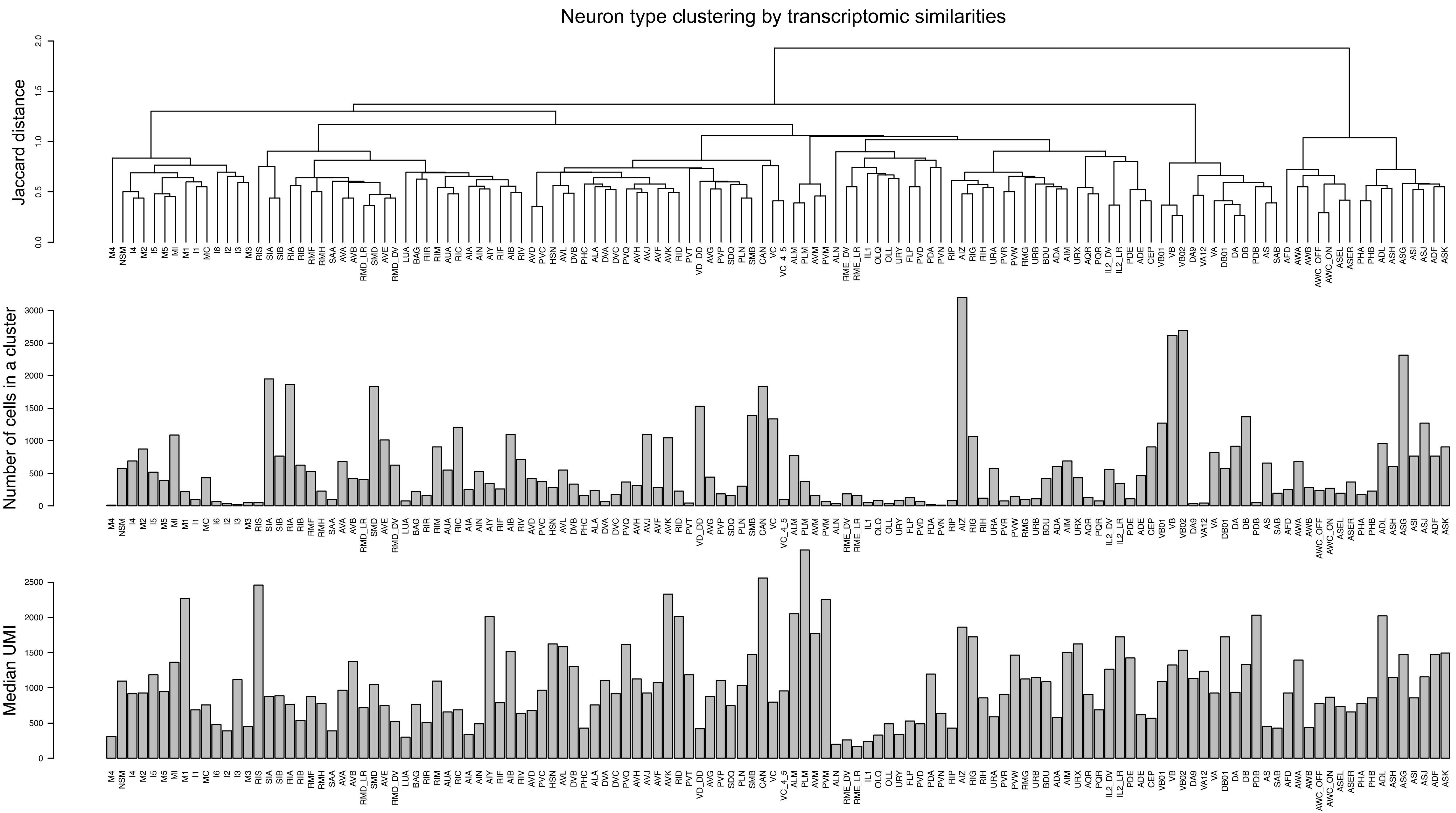


Supplementary Figure S2

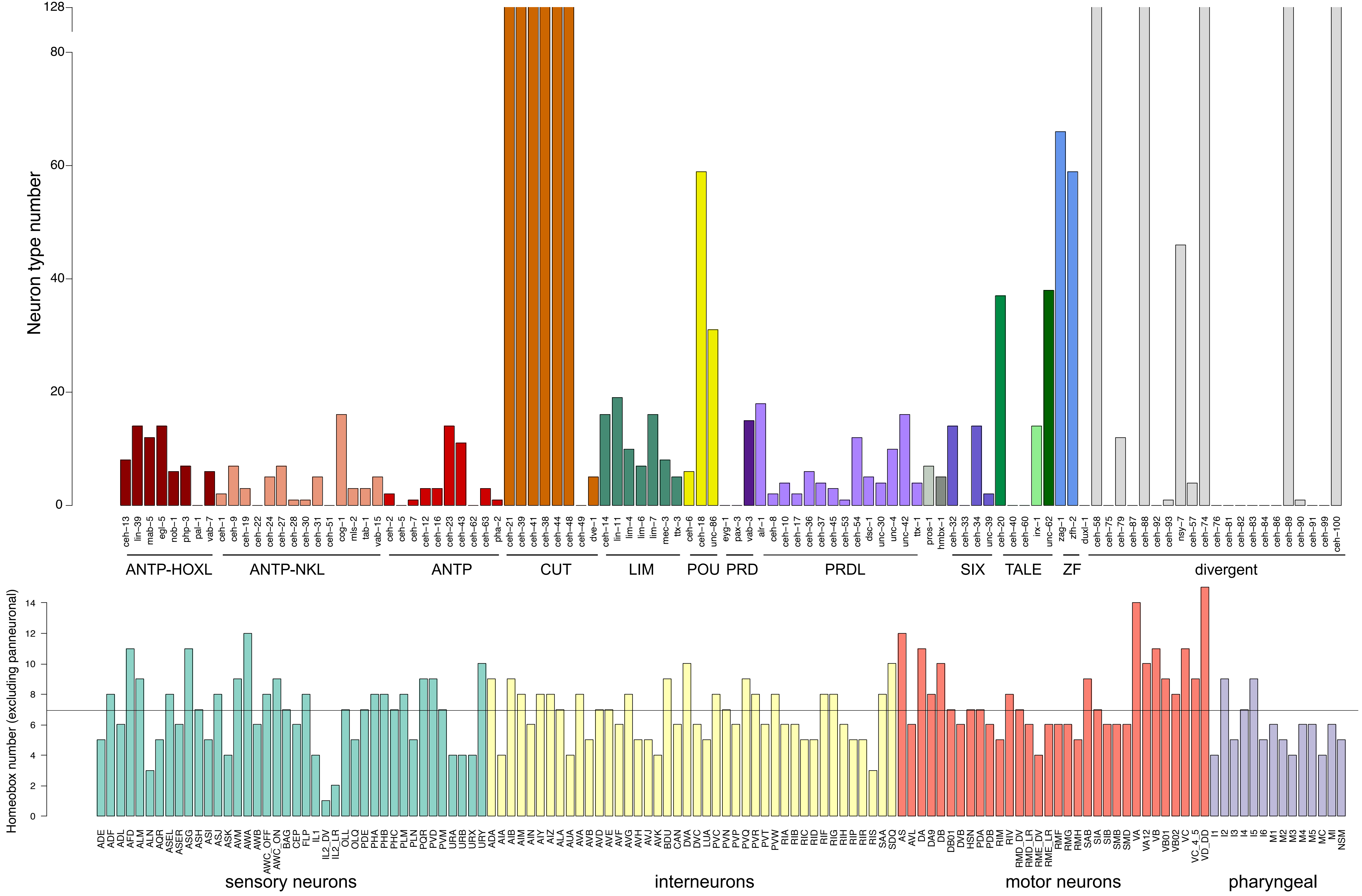


Supp. Figure S3

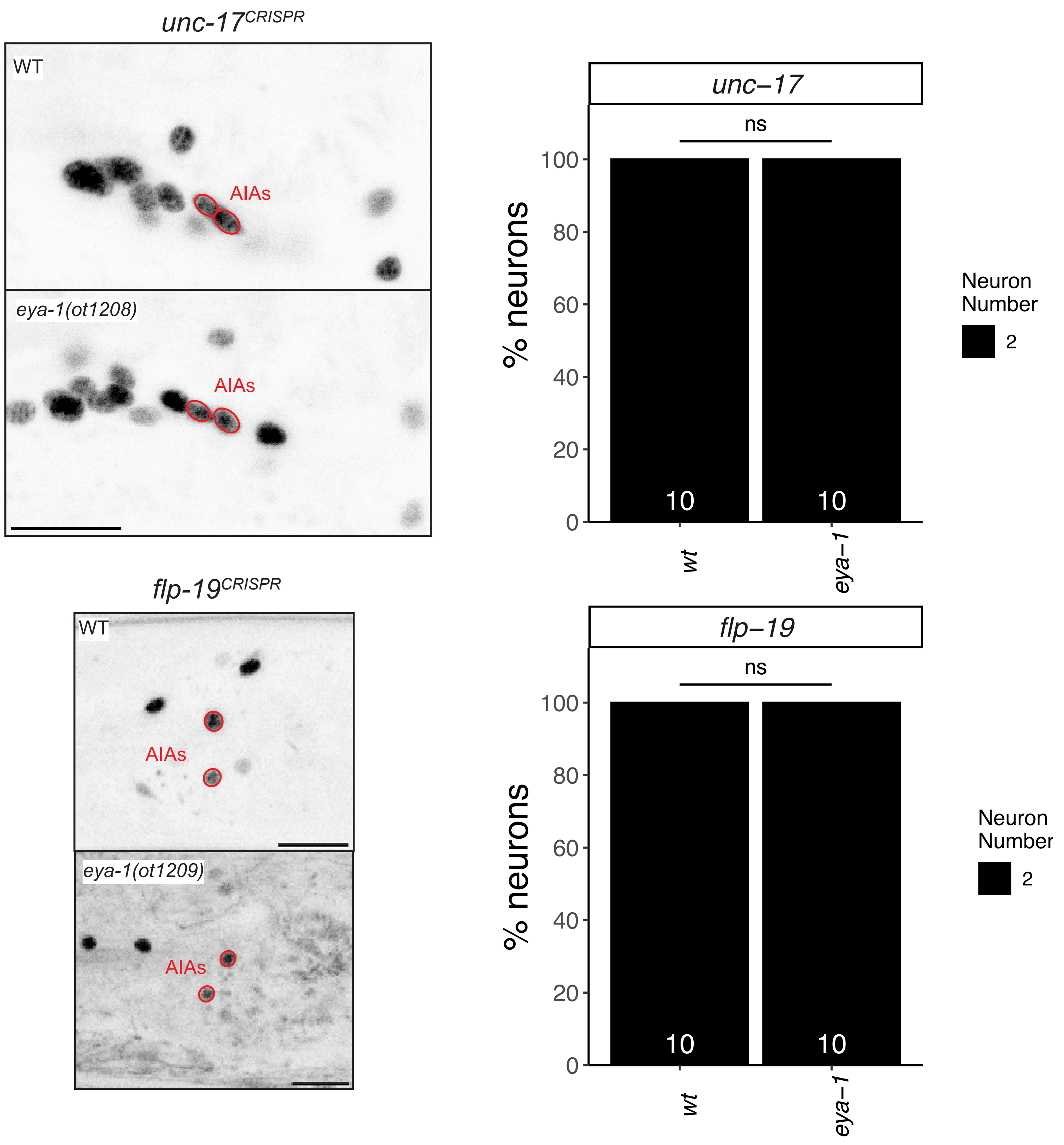


Supplementary Figure S4

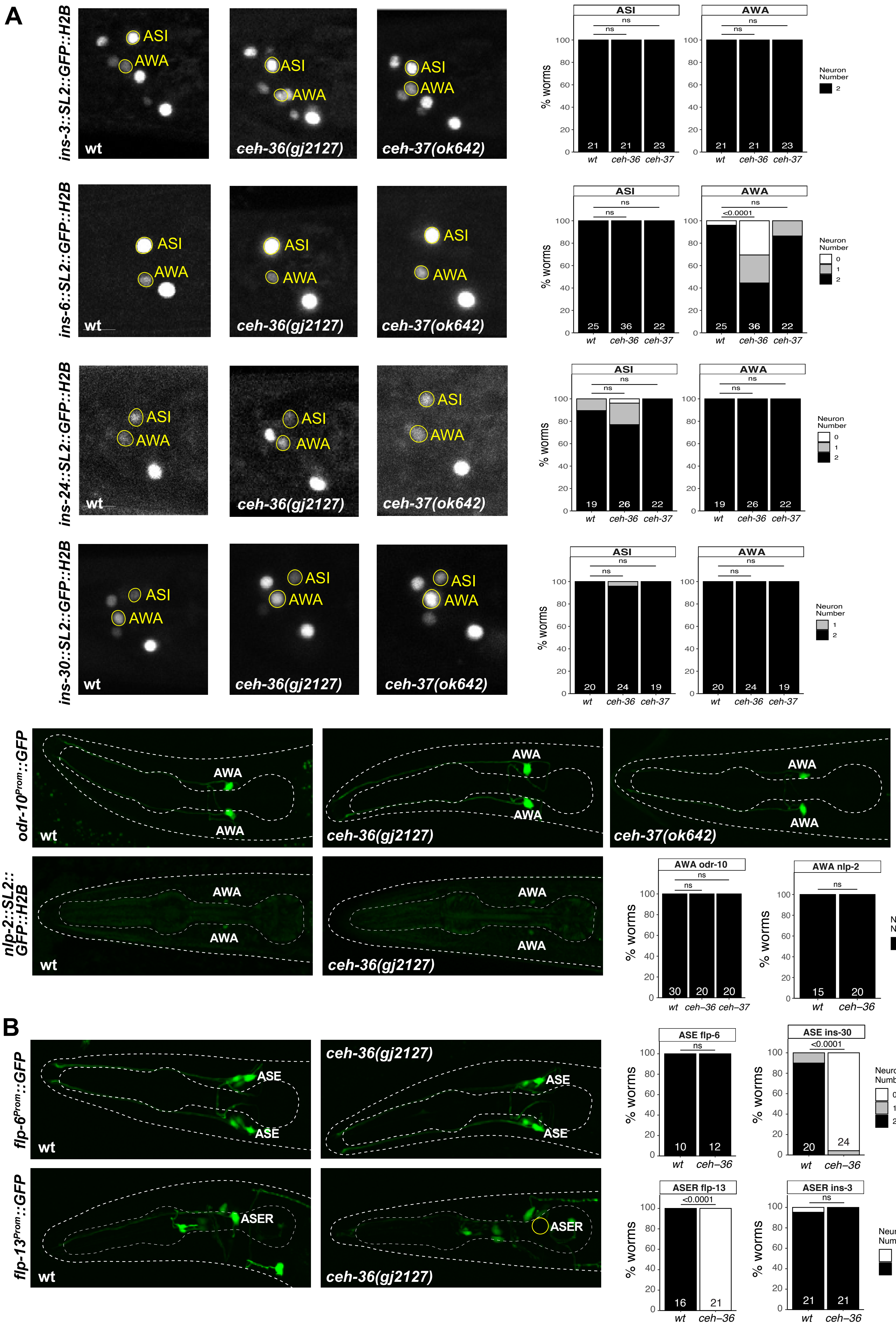


Supplementary Figure S5

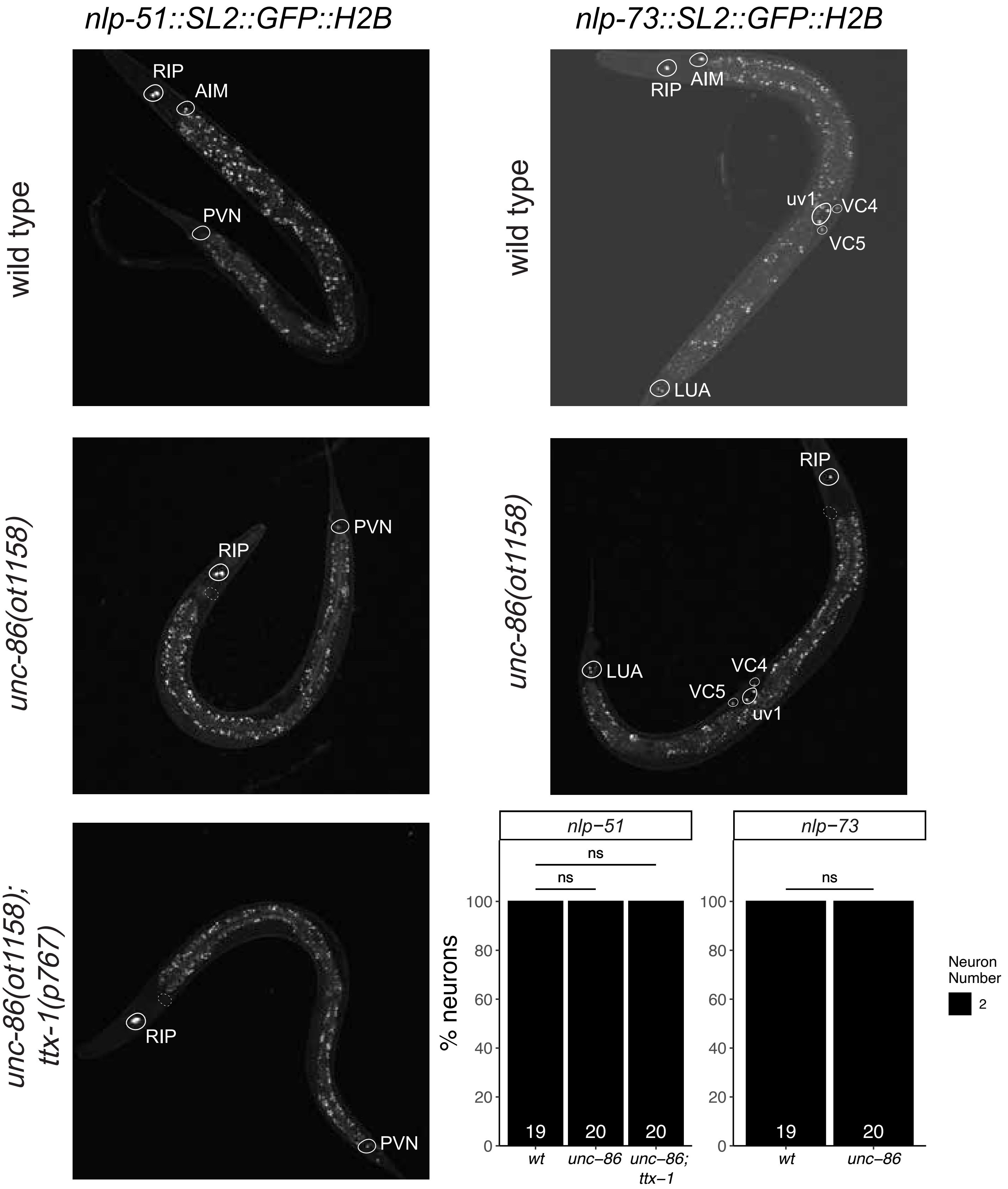


Suppl. Figure S6

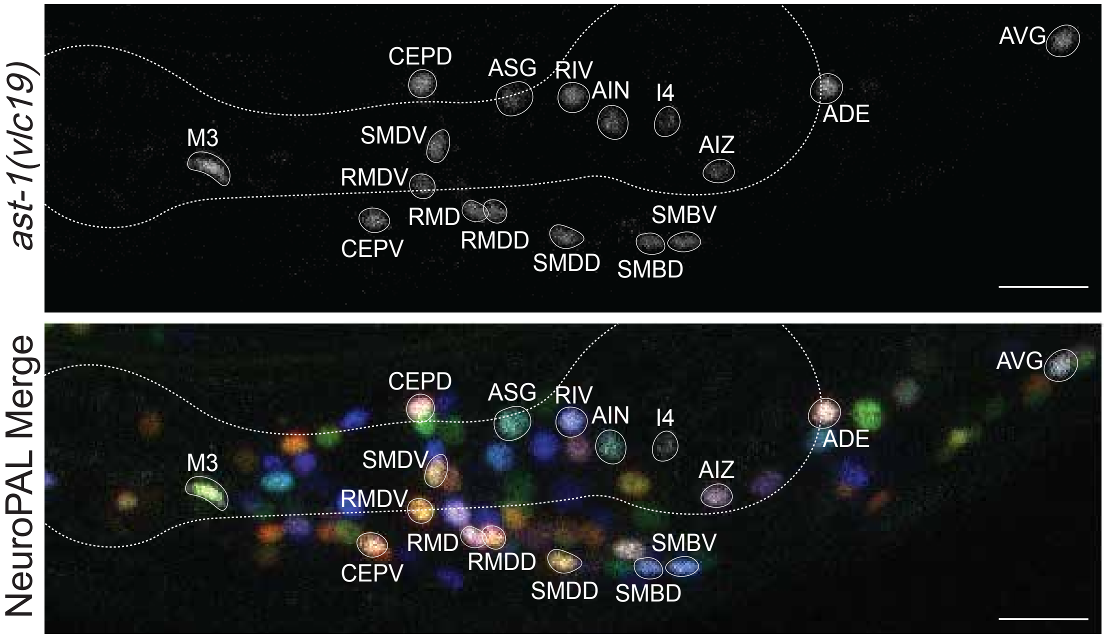


Suppl. Figure S7

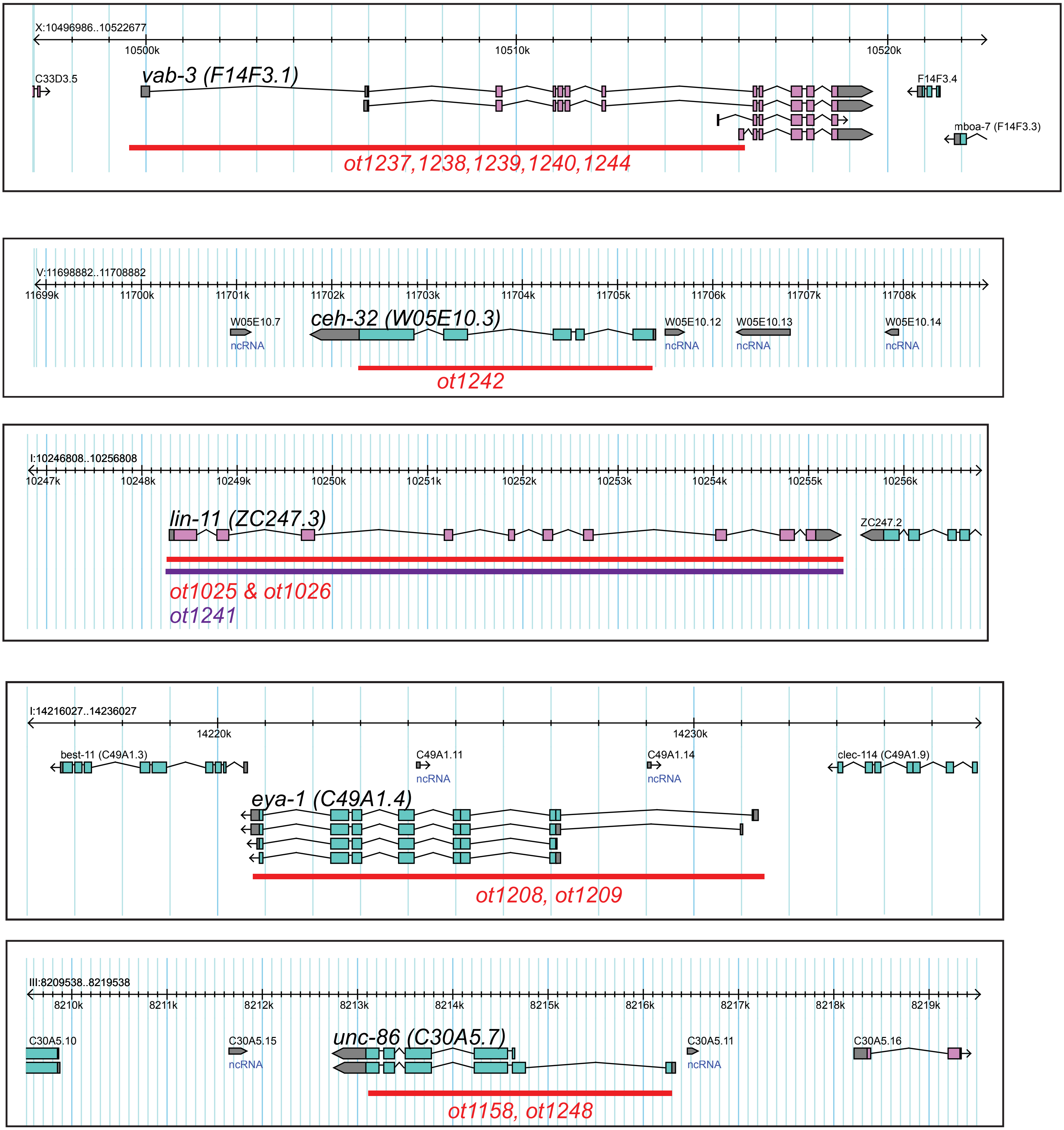


Supplementary Fig. S8
