## Supplementary material for "Widespread employment of conserved *C. elegans* homeobox genes in neuronal identity specification": Supp Table S3

**Table S3: Cells and identity markers not affected by homeobox mutant**

| Gene | Neuron class | Marker tested |
| --- | --- | --- |
| *lin-11* | DVA | NeuroPAL unaffected in *lin-11* null |
|  | RIC, RIM | *tbh-1* & *tdc-1* reporter unaffected in *lin-11* null |
| *unc-30* | ASG | NeuroPAL unaffected |
| *unc-62* | SDQ | NeuroPAL unaffected (but: only hypomorphic *unc-62* tested) |
|  | DVA | NeuroPAL unaffected (but: only hypomorphic *unc-62* tested) |
|  | PDB | NeuroPAL unaffected (but: only hypomorphic *unc-62* tested) |
|  | SAB | NeuroPAL unaffected (but: only hypomorphic *unc-62* tested) |
|  | RIM | NeuroPAL and *eat-4* unaffected (but: only hypomorphic *unc-62* tested) |
| *unc-86* | RIP | *nlp-51* and *nlp-73* unaffected |
| *ceh-9* | ASJ, AWB, SAA, RIV | *cho-1* fosmid transgene and unc-17 reporter allele unaffected |
|  | ADF | *cho-1* fosmid transgene and unc-17 reporter allele unaffected |
|  | PQR | *eat-4* fosmid unaffected |
| *ceh-19* | ADF | *cho-1* and *tph-1* fosmid transgenes unaffected |
|  | PHA | NeuroPAL and *eat-4* unaffected |
| *vab-3* | ADA | NeuroPAL, *nlp-82* reporter allele unaffected |
|  | ASK | *srg-8* (*otIs733*) unaffected |
| *ceh-32* | I2 | *gur-3 (nIs780), eat-4prom8 (otIs558*) unaffected |
|  | M1 | *unc-17prom3* transgene (*otIs661*) unaffected |
|  | ADL | *ift-20 srh-127 hlh-4* unaffected |
|  | RIB | *sto-3, unc-47, unc-46, cho-1* unaffected |
|  | RID | *flp-2 (ynIs57)* unaffected |
|  | RMH | *flp-32* reporter allele (*syb4374*) unaffected |
| *ceh-45* | RIB | *unc-47* fosmid unaffected |
| *lim-4* | SIB | *rig-6* unaffected; *unc-17* reporter allele slight dimming |
|  | SIA | NeuroPAL; *unc-17* unaffected |
|  | SMD | NeuroPAL; *unc-17* unaffected |
| *vab-7* | ADA, PHC | NeuroPAL; *eat-4* unaffected |
| *vab-15* | PVC | *hdIs32[glr-1::DsRed2]; ynIs54[flp-20p::GFP]* unaffected |
| *ceh-54* | I2, M3, BAG, AFD, AWC, ASE, ASG | *eat-4* unaffected |
| *ceh-57* | ADF, ASI, ASJ | NeuroPAL, *ins-24* unaffected |
| *ceh-31* | SDQ | *lad-2, unc-17, cho-1* unaffected |
|  | URA, URB | *unc-17, cho-1* unaffected |
| *pros-1* | DVA | *unc-17(syb4491)* and *nlp-12 (otIs706)* unaffected |
| *ceh-14* | ADA | NeuroPAL and *eat-4* reporter allele unaffected |
