## Supplementary material for "Widespread employment of conserved *C. elegans* homeobox genes in neuronal identity specification": Supp Table S5

**Supplementary Table S5: Strain list**

| **Strain Name** | **Genotype** | **DNA on array** | **Reference** |
| --- | --- | --- | --- |
| OH16745 | *vab-3(syb1653 ot1083) X* |  | This study |
| OH17961 | *vab-3(dev190([vab-3::mNeonGreen])* |  | [1] |
| PHX3426 | *ceh-27(syb2714[loxP] syb3286[loxP] syb3426[ceh27::GFP])* |  | This study |
| PHX5073 | *ceh-43(syb5073) [ceh-43::SL2::GFP::H2B]* |  | This study |
| PHX2880 | *ceh-16(syb2709[loxP] syb2880[ceh16::loxP::GFP])* |  | This study |
| PHX2934 | *ceh-36(syb2933[loxP]) ceh-36(syb2934[ceh36::loxP::GFP])* |  | This study |
| SYS629 | *ujIs113 (pie-1p::mCherry::H2B; nhr-2p::mCherry::his-24::let858 3’UTR; unc-119(+)) II; ceh-12 (devKi186 [ceh-12::mNeonGreen])* |  | [1] |
| SYS743 | *ujIs113 (pie-1p::mCherry::H2B; nhr-2p::mCherry::his-24::let858 3’UTR; unc-119(+)) II; unc-30 (hzhCR1 [unc-30::gfp])* |  | [1] |
| SYS608 | *ujIs113 (pie-1p::mCherry::H2B; nhr-2p::mCherry::his-24::let858 3’UTR; unc-119(+)) II; ceh-17 (devKi180 [mNeonGreen::ceh-17])* |  | [1] |
| SYS424 | *ujIs113 (pie-1p::mCherry::H2B; nhr-2p::mCherry::his-24::let858 3’UTR; unc-119(+)) II; ceh-5 (devKi103 [mNeonGreen::ceh-5])* |  | [1] |
| SYS397 | *ujIs113 (pie-1p::mCherry::H2B; nhr-2p::mCherry::his-24::let858 3’UTR; unc-119(+)) II; ceh-51 (devKi72 [mNeonGreen::ceh-51])* |  | [1] |
| SYS609 | *ujIs113 (pie-1p::mCherry::H2B; nhr-2p::mCherry::his-24::let858 3’UTR; unc-119(+)) II; ceh-82 (devKi181 [mNeonGreen::ceh-82])* |  | [1] |
| SYS635 | *ujIs113 (pie-1p::mCherry::H2B; nhr-2p::mCherry::his-24::let858 3’UTR; unc-119(+)) II; ceh-28 (devKi192 [mNeonGreen::ceh-28])* |  | [1] |
| SYS606 | *ujIs113 (pie-1p::mCherry::H2B; nhr-2p::mCherry::his-24::let858 3’UTR; unc-119(+)) II; ceh-2 (devKi17 [mNeonGreen::ceh-2])* |  | [1] |
| SYS634 | *ujIs113 (pie-1p::mCherry::H2B; nhr-2p::mCherry::his-24::let858 3’UTR; unc-119(+)) II; ceh-45 (devKi191 [mNeonGreen::ceh-45])* |  | [1] |
| SYS427 | *ujIs113 (pie-1p::mCherry::H2B; nhr-2p::mCherry::his-24::let858 3’UTR; unc-119(+)) II; ceh-10 (devKi101 [mNeonGreen::ceh-10])* |  | [1] |
| OH17991 | *otIs669; him-5 (e1490) V; ceh-31 (devKi250 [mNeonGreen::ceh-31]) X* | Bright NeuroPAL on V | [1] |
| MT19075 | *nIs352* | *eya-1p::gfp::eya-1* | [2] |
| OH17872 | *eya-1(ot1208) unc-17(syb4491)* |  | This study |
| OH17872 | *eya-1(ot1209) flp-19(syb3278)* |  | This study |
| PHX4678 | *ceh-30(syb4678[ceh-30::GFP])* |  | This study |
| OH17965 | *lin-11(ot1241);otIs711;; him-5* | *nlp-8prom::GFP; lin-15(+)* | This study |
| OH17870 | *lin-11(ot1026);unc-17(syb4491);otis669* |  | This study |
| OH17869 | *lin-11(ot1025); nIs107; zfIs1010* | *tdc1:mcherry; tbh-1::GFP* | This study |
| CB845 | *unc-30(e191)* |  | [3] |
|  | *egl-5(u202)* |  | [4] |
| LW227 | *mls-2(cc615)* |  | [5] |
| OH15655 | *otIs711* | *nlp-8prom::GFP; lin-15(+)* | [6] |
| OH13645 | *otIs518* | *eat-4fosmid::mcherry; him-5* | [7] |
| OH11124 | *otIs388* | *eat-4fosmid::sl2::yfp::H2B* | [7] |
| CX3260 | *kyIs37* | *odr-10::GFP; lin-15(+)* | [8] |
| OH17950 | *vab-3(ot1237)* |  | This study |
|  | *oxIs12* | *unc-47prom::GFP* | [9] |
| PHX1656 | *ceh-8(syb1656[ceh-8::GFP])* |  | [10] |
| OH1422 | *otIs138* | *ser-2(promoter3) + rol-6(su1006)* | [11] |
| OH16938 | *pha-1(e2123); otEx7697* | *des-2pOLQ::GFP + unc-122p::GFP + pha-1* | This study |
| OH17344 | *otIs849* | *ttll-9p(500 bp)::GFP + pha-1(+)* | This study |
| OH17345 | *otIs850* | *ttll-9p(500 bp)::GFP + pha-1(+)* | This study |
| CX5478 | *lin-15(n765); kyEx581* | *ocr-4::GFP + lin-15(+)* | [12] |
| OH12525 | *otIs521* | *eat-4prom8::tagRFP; ttx-3::gfp* | [13] |
| OH17951 | *vab-3(ot1238); otIs521* |  | This study |
| OH15034 | *otIs653* | *srg-8promASK cho-1promAIA GRASP* | [14] |
| OH17963 | *otIs879* | *sri-1prom::NLS::GFP pha-1rescue* | This study |
| OH17964 | *vab-3(ot1240); otIs879* |  | This study |
| OH17124 | *otIs837* | *otIs837 [unc-25(prom3)(del1)::GFP* | [15] |
| OH17970 | *ceh-32(ot1242)/+; otIs837* |  | This study |
| PHX4491 | *unc-17(syb4491[unc-17::T2A::GFP:H2B])* |  | [6] |
| PHX4257 | *eat-4(syb4257)[eat-4::T2A::GFP::H2B])* |  | [6] |
| OH17971 | *eat-4(syb4257)[eat-4::T2A::GFP::H2B]) vab-3(ot1244); otIs669* | Bright NeuroPAL on V | [6] |
| PHX4403 | *nlp-66(syb4403[nlp-66::SL2::gfp::H2B])* |  | This study |
| OH17952 | *nlp-66(syb4403[nlp-66::SL2::gfp::H2B])*  *vab-3(ot1239) X* |  | This study |
| OH15338 | *ceh-32(ok343); otEx7146* | *ceh-32fosmid WRM0637dA10 ;myo-2::RFP* | [16] |
| OH15601 | *myIs13; otIs703* | *[klp-6prom::GFP]* *[flp-3prom::mCherry]* | [16] |
| OH13026 | *otIs568* | *unc-46fosmid::SL2::H2B::mCHOPTI; pha-1 rescue* | [17] |
| PHX3278 | *flp-19(syb3278 [flp-19::T2A::3×NLS::GFP])* |  | [16] |
| *PHX5452* | *ins-1(syb5452[ins-1::SL2::gfp::H2B])* |  | [16] |
| PHX4514 | *dmsr-2(syb4514 [dmsr-2::SL2::gfp::H2B])* |  | [16] |
| OH17241 | *unc-86(ot1158)* |  | [16] |
| CB257 | *unc-39(e257)* |  | [3] |
| OH15363 | *him-5(e1490); otIs669* | Bright NeuroPAL on V | [18] |
| OH17491 | *him-5(e1490) unc-39(ot1173); otIs669* | Bright NeuroPAL on V | [16] |
| MT1859 | *unc-86(n846)* |  | [19] |
| OH17241 | *unc-86(ot1158)* |  | This study |
| CB644 | *unc-62(e644)* |  | [3] |
| NFB1369 | *ast-1(vlc19[ast-1::gfp])* |  | [20] |
| OH8251 | *otIs226* | *bas-1::gfp* | [21] |
| OH8249 | *otIs224* | *cat-1::gfp* | [22] |
|  | *nIs118* | *cat-2::gfp* | [23] |
| OH8250 | *otIs225* | *cat-4::gfp* | [22] |
| BY200 | *vtIs1* | *dat-1::gfp* | [24] |
| PHX3195 | *flp-33(syb3195[flp-33::T2A::3xNLS::GFP])* |  | This study |
| OH7197 | *ast-1(ot417); otIs199[cat-2::gfp; rgef-1::dsred]* |  | [21] |
| OH7323 | *ceh-43(ot406); vtIs1; vsIs33* |  | [22] |
| OH18009 | *unc-86(ot1248); ceh-43(ot406); vtIs1; vsIs33* |  | This study |
| PHX4257 | *eat-4(syb4257[eat-4::T2A::GFP::H2B])* |  | [25] |
| OH15422 | *ceh-14(ot900)* |  | [26] |
| OH10210 | *otEx4530* | *cho-1prom3::DsRed2* | [13] |
| OH10765 | *otIs358* | *ser-2prom2::gfp; pha-1(e2123)* | [27] |
| CX3300 | *kyIs51* | *odr-2prom::gfp* | [28] |
| OP756 | *unc-119(tm4063) III; wgIs756 [ceh-13::TY1::EGFP::3XFLAG + unc-119(+)]* | *ceh-13* fosmid reporter | [29] |
| PHX3323 | *flp-14(syb3323[flp-14::T2A::3xNLS::GFP])* |  | [16] |
| PHX3212 | *flp-21(syb3212[flp-21::T2A::3xNLS::GFP])* |  | This study |
| PHX4513 | *flp-5(syb4513[flp-5::SL2::GFP::H2B])* |  | [16] |
| FR431 | *ceh-13(sw1)/qC1 dpy-19(e1259) glp-1(q339) III* |  | [30] |
| KRA758 | *ceh-13(sw1)/hT2 III; flp-21(syb3212 [flp-21::T2A::3×NLS::GFP]) V* |  | This study |
| KRA759 | *mab-5(e1239) III; flp-6 (syb3203 [flp-6::T2A::3xNLS::GFP])V* |  | This study |
| KRA760 | *mab-5(e1239) III; nlp-40(syb3208 [nlp-40::T2A::3xNLS::GFP])I* |  | This study |
| KRA761 | *mab-5(e1239) III; otIs92[flp-10::gfp]V* |  | This study |
| OH17006 | *otIs576; otIs669; him-5(e1490) V* | *otIs576* [*unc-17 fosmid::GFP; lin-44::YFP*]  *otIs669:* NeuroPAL | This study |
| OH17049 | *tab-1(ok2198) II; otIs576; otIs669; him-5(e1490) V* | *otIs576* [*unc-17 fosmid::GFP; lin-44::YFP*]  o*tIs669*: NeuroPAL | This study |
| OH17012 | *nlp-42(syb3238[nlp-42::T2A::3xNLS::GFP]); otIs669; him-5(e1490) V* |  | This study |
| OH17068 | *tab-1(ok2198) II; nlp-42(syb3238); otIs669 him-5(e1490) V* |  | This study |
| OH17037 | *otIs576; otIs669; him-5(e1490) V; ttx-3(ot22) X* |  | This study |
| OH17053 | *otIs669 him-5(e1490) V; ttx-3(ot22) X; nlp-42 (syb3238)* |  | This study |
| NY2050 | *ynIs50 (pflp-22::GFP); him-5(e1490)* |  | [31] |
| OH17973 | *ynIs50 (pflp-22::GFP); ceh-14(ot900)* |  | [31] |
| PHX4413 | *flp-27(syb4413 [flp-27::SL2::GFP::H2B])* |  | This study |
| PHX3212 | *flp-21(syb3212 [flp-21::T2A::3×NLS::GFP])* |  | This study |
| PHX3411 | *nlp-13 (syb3411 [nlp-13::T2A::3XNLS::GFP])* |  | This study |
| OH17974 | *flp-27(syb4413 [flp-27::SL2::GFP::H2B]); ceh-14(ot900)* |  | This study |
| OH17975 | *flp-21(syb3212 [flp-21::T2A::3×NLS::GFP]); ceh-14(ot900)* |  | This study |
| OH17976 | *nlp-13 (syb3411 [nlp-13::T2A::3XNLS::GFP]); ceh-14(ot900)* |  | This study |
| PHX5421 | *ins-3(syb5421[ins-3::SL2::gfp::H2B]) II.* |  | This study |
| PHX5463 | *ins-6(syb5463[ins-6::SL2::gfp::H2B]) II.* |  | This study |
| PHX5447 | *ins-24(syb5447[ins-24::SL2::gfp::H2B]) I* |  | This study |
| PHX5526 | *ins-30(syb5526[ins-30::SL2::gfp::H2B]) I* |  | This study |
| PHX5697 | *nlp-2(syb5697 [nlp-2::SL2::GFP::H2B]) X* |  | This study |
| NY2067 | *ynIs67 III; him-5(e1490) V.* | *flp-6prom::GFP* | [31] |
| NY2037 | *ynIs37* | *flp-13prom::GFP* | [31] |
| OH17795 | *ceh-36(gj2127) X* |  | [32] |
| RB823 | *ceh-37(ok642) X* |  | [33] |
| PHX2805 | *nlp-51::SL2::GFP::H2B(syb2805)* |  | This study |
| PHX4406 | *nlp-73::SL2::GFP::H2B(syb4406)* |  | This study |
| OH17968 | *nlp-51::SL2::GFP::H2B(syb2805); unc-86(ot1158)* |  | This study |
| OH17992 | *unc-86(ot1158); nlp-73::SL2::GFP::H2B(syb4406)* |  | This study |
| OH17957 | *nlp-51::SL2::GFP(syb3936); unc-86(ot1158); ttx-1(p767)* |  | This study |
| OH16366 | *otEx7603* | *nlp-52p::gfp, pha-1(+)* | This study |
